# A diabetic *milieu* increases cellular susceptibility to SARS-CoV-2 infections in engineered human kidney organoids and diabetic patients

**DOI:** 10.1101/2021.08.13.456228

**Authors:** Elena Garreta, Patricia Prado, Megan Stanifer, Vanessa Monteil, Carmen Hurtado del Pozo, Asier Ullate-Agote, Amaia Vilas-Zornoza, Juan Pablo Romero, Gustav Jonsson, Roger Oria, Alexandra Leopoldi, Astrid Hagelkruys, Daniel Moya-Rull, Federico González, Andrés Marco, Carolina Tarantino, Pere Domingo-Pedrol, Omar HasanAli, Pedro Ventura-Aguiar, Josep María Campistol, Felipe Prosper, Ali Mirazimi, Steeve Boulant, Josef M. Penninger, Nuria Montserrat

## Abstract

SARS-CoV-2 infections lead to a high risk of hospitalization and mortality in diabetic patients. Why diabetic individuals are more prone to develop severe COVID-19 remains unclear. Here, we established a novel human kidney organoid model that mimics early hallmarks of diabetic nephropathy. High oscillatory glucose exposure resulted in metabolic changes, expansion of extracellular membrane components, gene expression changes determined by scRNAseq, and marked upregulation of angiotensin-converting enzyme 2 (ACE2). Upon SARS-CoV-2 infection, hyperglycemic conditions lead to markedly higher viral loads in kidney organoids compared to normoglycemia. Genetic deletion of ACE2, but not of the candidate receptor BSG/CD147, in kidney organoids demonstrated the essential role of ACE2 in SARS-CoV-2 infections and completely prevented SARS-CoV-2 infection in the diabetogenic microenvironment. These data introduce a novel organoid model for diabetic kidney disease and show that diabetic-induced ACE2 licenses the diabetic kidney to enhanced SARS-CoV-2 replication.

## Introduction

Coronavirus disease 2019 (COVID-19) is an infectious disease caused by SARS-Coronavirus 2 (SARS-CoV-2). COVID-19 patients display influenza-like symptoms ranging from mild disease to severe lung injury. A high percentage of severe COVID-19 patients display symptoms in other organs, most notably the gastrointestinal tract, cardiovascular system and the kidney. Several conditions have been linked to the risk of developing severe COVID-19, including genetic predisposition [1–3], immune related-responses [4, 5] obesity, or Diabetes mellitus (DM) [6]. Both COVID-19 and DM are associated with acute and chronic inflammation and both disease conditions can impact each other in terms of clinical progression and disease outcome [7]. Notably, SARS-CoV-2 infections lead to acute kidney injury in >20% of hospitalized patients [8] and higher rates of mortality have been reported in patients with pre-existing DM [9].

Human organoids are excellent model systems to assess the pathogenesis of SARS-CoV-2 infection and to identify and validate potential drug targets. We have previously shown that both kidney and vascular organoids derived from human pluripotent stem cells (hPSCs) support SARS-CoV-2 infections which was blocked in the presence of clinical grade human recombinant soluble Angiotensin Converting Enzyme 2 (ACE2) [10]. However, besides the great utility of organoids in SARS-CoV-2 research, organoids have not yet been developed that model human co-morbidities associated with severe COVID-19, such as DM.

Here we show that high glucose oscillations in engineered human kidney organoids led to phenotypic, transcriptional, and metabolic alterations reminiscent to diabetic kidney tissue in human patients. These diabetic conditions enhanced ACE2 expression and SARS-CoV-2 infection, which was validated in human proximal tubular cells isolated from diabetic kidney biopsies. Genetic deletion of ACE2, but not BSG/CD147, another candidate receptor for Spike binding, completely abrogated SARS-CoV-2 infections under normal and diabetic conditions. This study provides mechanistic evidence on metabolic alterations that can increase cellular susceptibility to SARS-CoV-2 infections and unequivocally establishes ACE2 as the critical SARS-CoV-2 receptor in the human kidney, even under diabetic conditions.

## Results

### Establishment of diabetic human kidney organoids

To study the impact of hyperglycemia on renal cells in SARS-CoV-2 infections, we established a novel procedure to generate diabetic-like kidney organoids from hPSCs. Adapting our previous protocol [10] we first induced posterior primitive streak (PPS) fate by exposing monolayer hPSCs (day 0) to the Wnt agonist CHIR (8 μM) for 3 days. Then PPS committed cells were cultured in FGF9 (200 ng/mL), activin-A (10 ng/mL) and heparin (1μg/mL) for 24h to promote posterior intermediate mesoderm (PIM). On day 4, PIM-committed progenitors were treated with CHIR (5 μM), FGF9 (200 ng/mL) and heparin (1μg/mL) for 1h, and subsequently aggregated into spheroids containing nephron progenitor cells (NPCs) (day 8). In the presence of FGF9 (200 ng/mL), NPC-spheroids formed self-organized renal vesicle (RV) structures (day 11). From that stage RVs autonomously developed into free floating segmented nephron-like structures until day 16 (nephrogenesis) (Figure 1A).

**Figure 1.**
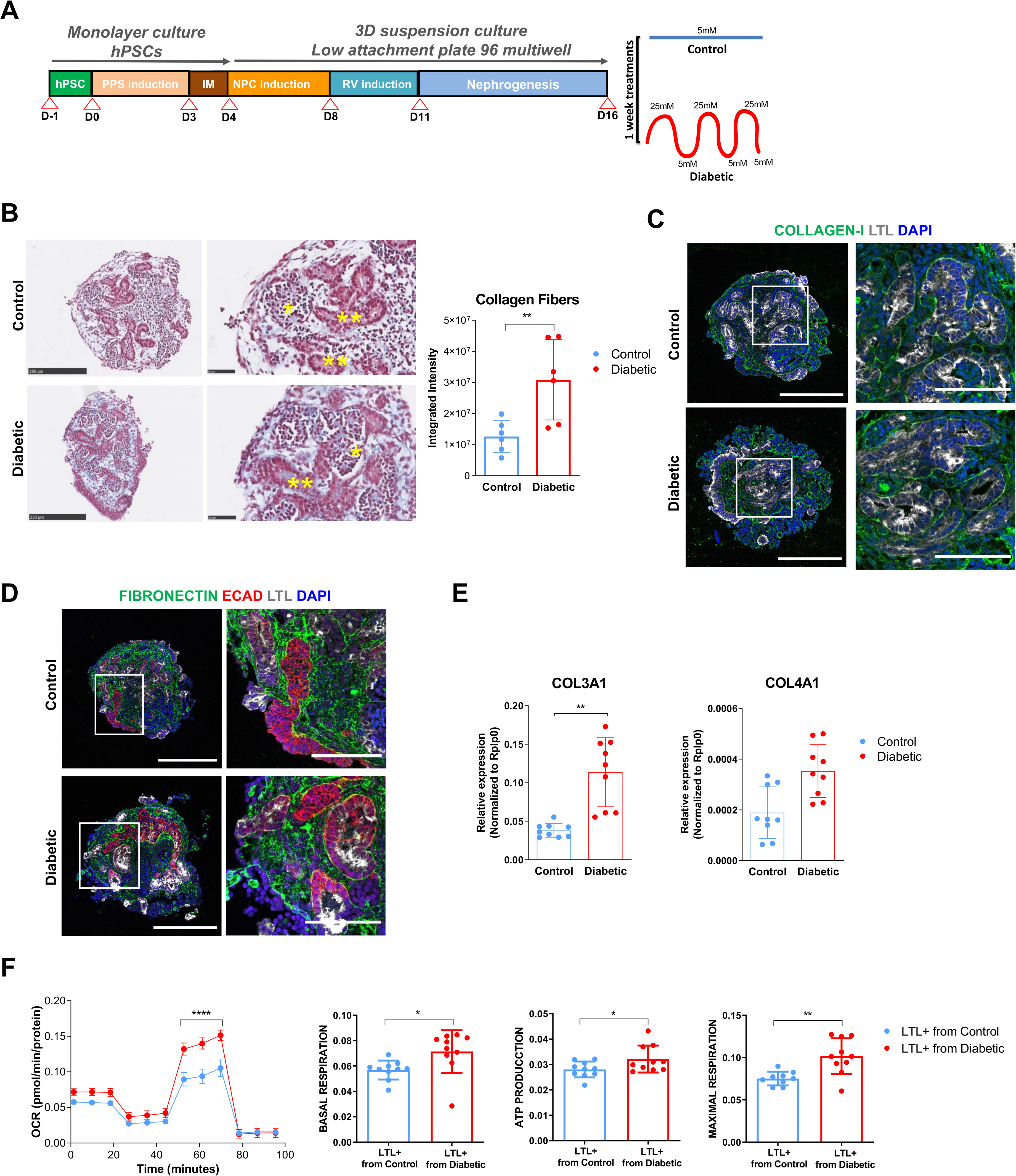
High Oscillatory Glucose Conditions Induce Early Hallmarks of the Diabetic Kidney Disease in Human Kidney Organoids. A. Experimental scheme for the generation of human kidney organoids from hPSCs. Briefly, hPSCs are induced to a first phase of differentiation into posterior primitive streak (PPS) fate and differentiated into intermediate mesoderm (IM) fate as monolayer cell cultures. At that day of differentiation (Day 4; D4) IM-like cells are aggregated and further differentiated in three-dimensional (3D) culture conditions. From that stage 3D cultures are further differentiated into renal vesicle (RV) stage which transition autonomously into nephron stage organoids in the absence of growth factors. After 16 days in suspension kidney organoids are exposed into 5 mM glucose (Control) or high oscillatory glucose (Diabetic) conditions changing media every 24 hours in the presence of 5 to 25 mM glucose. B. Trichrome Masson staining of kidney organoids treated for 7 days in Control or Diabetic conditions. Magnified views of glomerular (*) and tubular (**) structures are shown. Scale bars, 250 μm, 50 μm (magnified views). Corresponding quantification of collagen fibers in the Trichrome Masson images. Y-axis represents integrated intensity. The data are presented as mean ± SEM. N = 6 organoids/group. C. Representative immunofluorescence staining of COLLAGEN-I (green), LTL (grey) and DAPI (blue) in kidney organoids in Control and Diabetic conditions. Scale bars, 250 μm, 100 μm (magnified views). D. Representative immunofluorescence staining of FIBRONECTIN (green), E-CADHERIN (ECAD; red), LTL (grey) and DAPI (blue) in kidney organoids exposed to Control or Diabetic conditions for 7 days. Scale bars, 250 μm, 100 μm (magnified views). E. mRNA expression level of ECM markers (COL3A1, COL4A1) in kidney organoids exposed to Control or Diabetic conditions for 7 days. The data are represented as mean ± SEM. N>9 independent experimental replicates from a pool of 12 organoids/group; *P < 0.05, **P < 0.01 ***P < 0.001, paired Student’s t-test. F. Kinetic oxygen consumption rate (OCR) response, basal respiration and spare respiratory capacity, cellular ATP production and inner mitochondrial membrane proton leak in LTL+ cells isolated from control and diabetic kidney organoids. Data are normalized to total protein. Mean ± SEM is calculated from 3 biological replicates/group performing 3 technical replicates each. **p<0.01; ***p<0.001***, two-way ANOVA, followed by Bonferroni post-test. See also Figure S1; Figure S2; Extended Data 1; Extended Data 2; Extended Data 3.

Diabetic patients exhibit oscillatory levels in glucose which is thought to be important for driving DM-associated pathologies. Indeed, the term “metabolic memory” coins the pathogenic alterations induced by hyperglycemia long after accomplishment of glycemic control [11, 12]. To emulate *in vitro* diabetic patient-like oscillations in glucose levels, we determined the optimal glucose concentrations and time points for “diabetic” inventions. Following multiple pilot experiments, we delineated a culture set up using continuous low glucose (5 mM, termed control) or high glucose in an oscillatory fashion (alternating 5 to 25 mM every 24 hours, termed “diabetic” conditions) (Figure 1A). Kidney organoids under both control and diabetic culture conditions showed the presence of glomerular-like and renal tubular-like structures to a similar extent, as determined by immunodetection of PODXL^+^ podocyte-like cells and Lotus Tetraglobus Lectin (LTL)^+^ proximal tubule cells (Figure S1A) and qPCR analysis for the proximal tubule marker SLC12A1, SLC3A1, SLC12A3, CDH16 and the podocyte marker genes MAFB, PODXL, and WT1 (Figure S1B). Transmission electron microscopy (TEM) showed that tubular-like cells from both control and diabetic kidney organoids exhibited prominent brush borders (bb) and high mitochondrial (mt) content; podocyte-like cells presented typical features of native developing podocytes including apical microvilli and primary (pp) and secondary (sp) cell processes adjacent to the basement membrane (bm) (Figure S1C).

Changes in kidney extracellular matrix proteins are associated with chronic kidney disease (CKD) and renal fibrosis in diabetes [13, 14]. Importantly, high oscillatory glucose treatment resulted in a marked increase in collagen deposition in kidney organoids compared to low glucose controls (Figure 1B,C, Figure S1D; Extended Data 1A,B). Similarly, fibronectin deposits were detected in the tubulointerstitial areas within kidney organoids (Figure 1D; Extended Data 2). We also observed an upregulation of collagen III and collagen IV expression by qPCR (Figure 1E). PAS staining and immunohistochemistry to identify the basement membrane proteins LAMININ and Collagen IV in combination with the podocyte marker NEPHRIN and proximal tubule marker LTL showed that the composition and integrity of basement membranes was largely preserved under diabetic conditions (Figure S1E-F; Extended Data 3A,B). The glycolysis-associated genes LDHA and HK2 as well as CD36, a multifunctional receptor with roles in lipid accumulation, inflammatory signaling, and kidney fibrosis [15], were significantly upregulated in the diabetic kidney organoids, whereas PGC1α (a key regulator of mitochondrial biogenesis involved in diabetic nephropathy [16]) was downregulated (Figure S1G).

To assess whether the diabetic milieu has altered cellular metabolism, we isolated proximal tubular-like cells by fluorescent-activated cell sorting of the LTL^+^ cell fraction (Figure S2A) accounting for 9.9 ± 2.1 % (mean ± SD, n = 3) and 8.2 ± 1.1 % (mean ± SD, n = 3) of the total live cells from control and diabetic kidney organoids (Figure S2B). LTL^+^ cells were plated until passage 4 under 5 mM glucose conditions for 2 months. PGC1α expression in LTL^+^ cells was significantly decreased in cells isolated from diabetic organoids as shown by qPCR (Figure S2C) and immunofluorescence (Figure S2D). The expression of the tubular markers LTL and Na-K ATPase were unchanged (Figure S2D). The reduction in PGC1α levels were concomitant with increases in the maximum oxygen consumption rate (OCR), a measurement of mitochondrial respiration, basal and maximal respiration, as well ATP synthesis (Figure 1F). Of note, our results are in line with increased mitochondrial fitness in proximal tubular cells assessed by OCR measurement at early phases of diabetes in animal models (1–4 weeks after induction of diabetes) [17, 18]. These data show that exposure of human kidney organoids to high oscillatory glucose leads to transcriptional changes, fibrotic alterations, and metabolic mitochondrial rewiring in tubular cells, early hallmarks of diabetic kidney disease. Our culture system also allowed for the generation of LTL^+^ cells recapitulating hallmarks of an early diabetogenic-like phenotype after removal of the diabetogenic insult, indicative of metabolic memory.

### Diabetic conditions induce ACE2 expression in human kidney organoids

The Angiotensin-converting enzyme 2 (ACE2) has been previously identified as a key host cell-surface receptor sufficient for SARS-CoV-2 [19–21]. Of note, ACE2 has also been previously shown to control the progression of CDK in multiple animal models [22]. Although it has been reported that glucose can induce ACE2 expression in cell lines [23], it is still controversial whether DM results in up or downregulation of ACE2 [24–26], making it paramount to test the effects of glucose on ACE2 expression in complex human tissue-like cultures as organoids. Our analysis indicated that in the kidney organoids, we detected ACE2 expressing cells predominantly in LTL^+^ proximal tubule cells as shown by immunofluorescence (Figure 2A; Extended Data 4), thereby recapitulating ACE2 expression in the native human and mouse kidney [27, 28]. Importantly, the oscillatory glucose treatment promoted a significant upregulation of ACE2 expression compared to control conditions at the protein and mRNA levels (Figure 2 B,C; Extended Data 4). Since mRNA stability plays a major role in gene expression regulation, we hypothesized that hyperglycemia might affect in the half-life of ACE2 mRNA. To test this idea, kidney organoids were challenged with the RNA transcription inhibitor Actinomycin-D at 0, 2, 4 and 8 hours. Indeed, oscillatory glucose conditions led to an increase of ACE2 mRNA stability compared to control culture conditions (Figure 2D). Thus, a diabetic milieu results in marked upregulation of ACE2 in proximal tubular-like cells in human kidney organoids.

**Figure 2.**
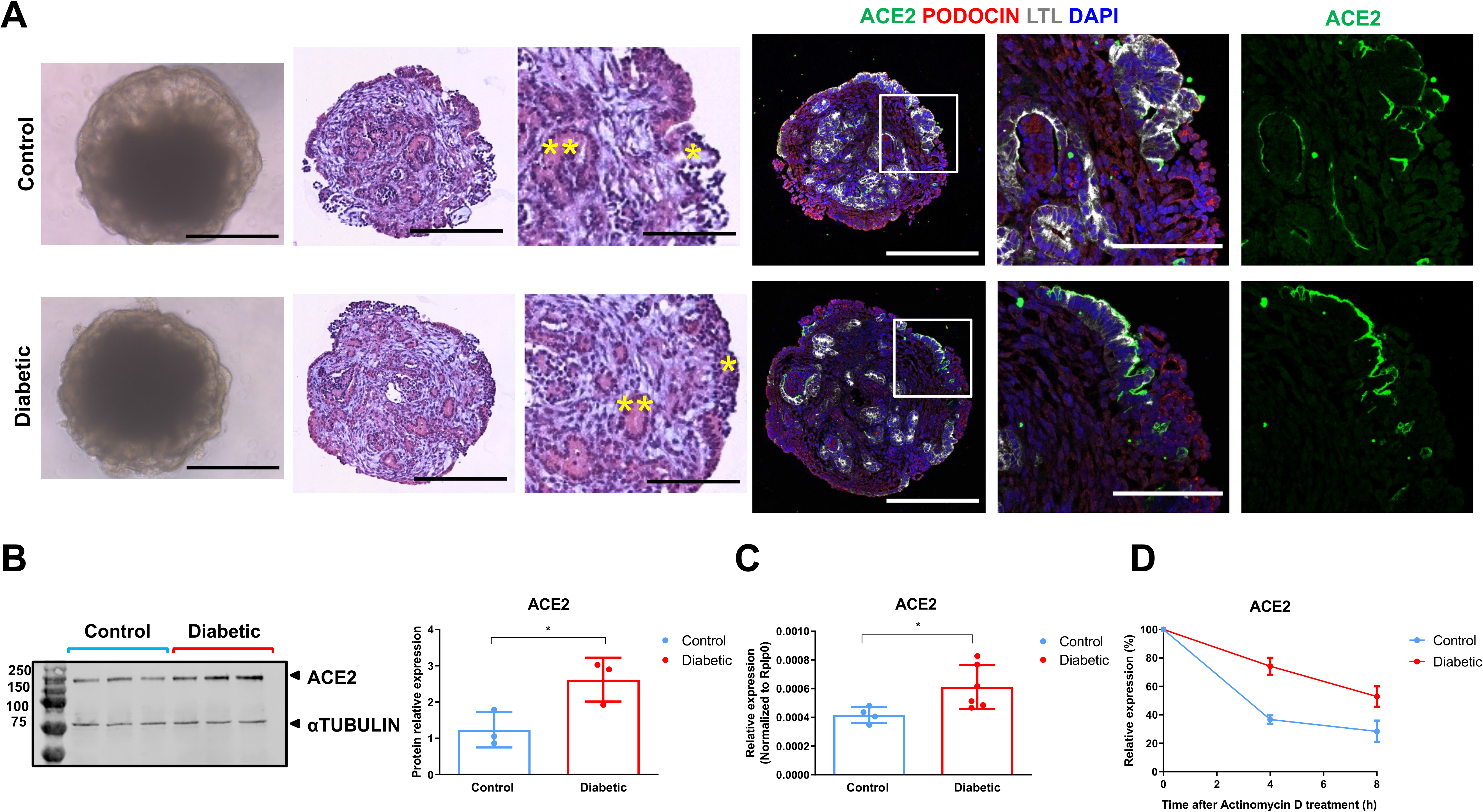
High Oscillatory Glucose Conditions Induce ACE2 Expression in Human Kidney Organoids. A. Representative bright-field images of kidney organoids exposed to Control or Diabetic conditions for 7 days. Representative Hematoxylin and Eosin staining. Asterisks highlight podocyte-like cells (*) or tubular -like structures (**). Scale bars, 250 μm, 100 μm (magnified views). Consecutive sections were stained for ACE2 (green), PODOCIN (red) and using Lotus Tetraglobus Lectin (LTL, tubular cells marker, grey). Nuclei were counterstained with DAPI (blue). Scale bars, 250 μm, 100 μm (magnified views). B. Protein levels of ACE2 in kidney organoids exposed to Control or Diabetic conditions for 7 days are shown by Western Blot. aTUBULIN was used as loading control. The quantification of changes in the protein expression for ACE2 is shown below as data represented as mean ± SEM. *n* = 3 independent experimental replicates from a pull of 12 organoids/group; \**P* < 0.05, paired Student’s *t*-test. C. mRNA expression level of ACE2 in kidney organoids exposed to Control or Diabetic conditions for 7 days. The data are represented as mean ± SEM. *n* = 5 independent experimental replicates from a pool of 12 organoids/group; \**P* < 0.05, paired Student’s *t*-test. D. ACE2 mRNA expression levels in kidney organoids exposed to Control or Diabetic conditions for 7 days treated with actinomycin D (1 mg/ml) for 30 min and then cultured for 2, 4, 6 and 8 hours. See also Extended Data 4.

### Increased SARS-CoV-2 infection in diabetic kidney organoids

Our results so far showed that a high oscillatory glucose treatment reproduced early hallmarks of the *in vivo* diabetic kidney and such treatment led to upregulation of ACE2. Therefore we aimed to evaluate the impact of these “diabetic” changes on SARS-Co-V-2 infections. Diabetic and control kidney organoids were infected with SARS-CoV-2, recovered one day post-infection (1dpi) (Figure 3A), and analyzed using whole mount immunofluorescence (Figure 3B). Control organoids were productively infected as detected by immunostaining for virus nucleoprotein (NP) (Figure 3B,C; Figure S3). Infected cells within organoids were primarily ACE2^+^ and LTL^+^ proximal tubular-like cells (Figure 3B). Remarkably, organoids exposed to the diabetic milieu showed significantly enhanced SARS-CoV-2 infections as quantified by the percentage of NP^+^ cells in the whole organoid by confocal microscopy (Figure 3B,C). Enhanced SARS-CoV-2 infections in diabetic kidney organoids were confirmed using qPCR to detect viral RNA (Figure 3D).

**Figure 3.**
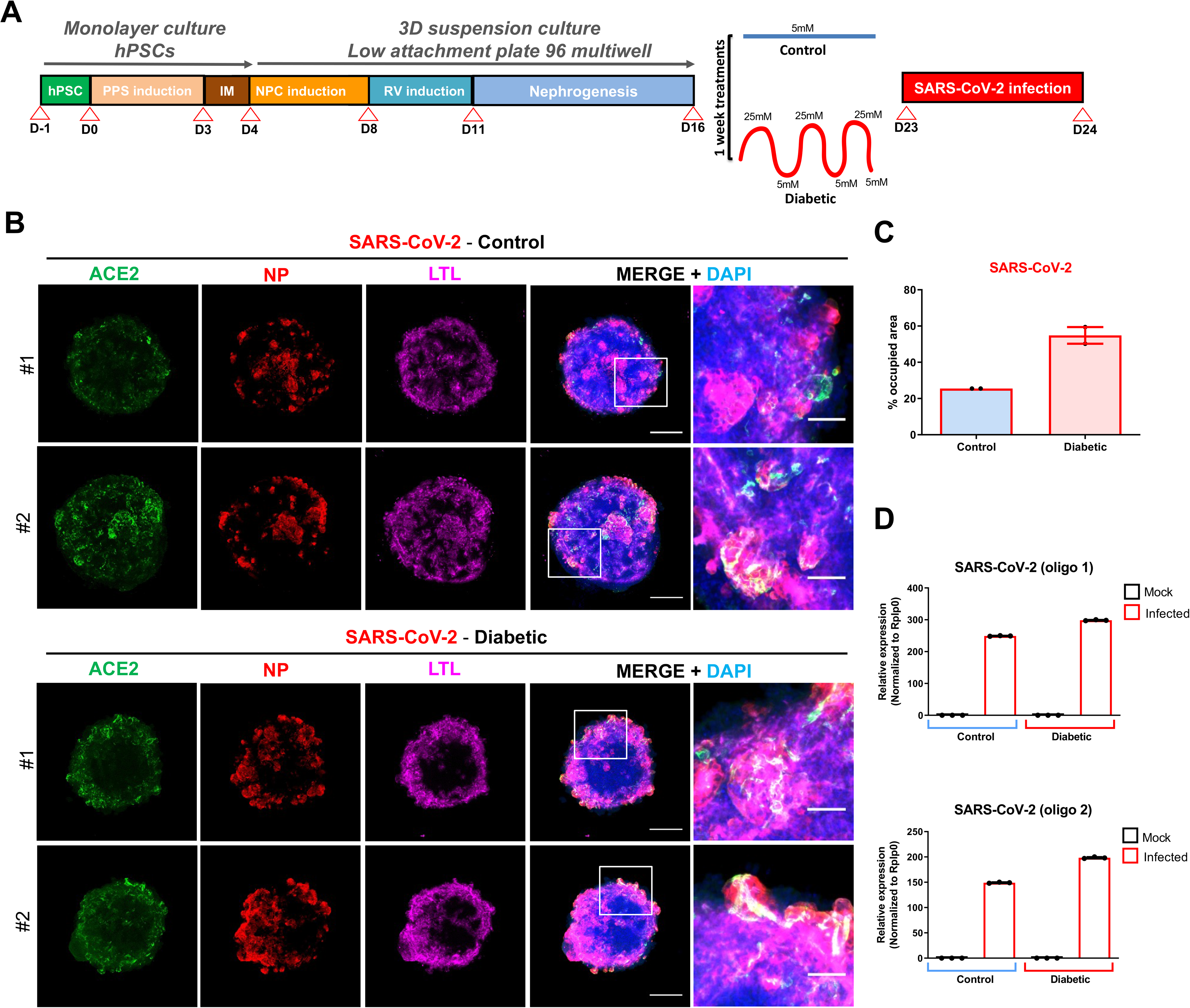
A Diabetic *Milieu* Promotes Enhanced SARS-CoV-2 Infections in Human Kidney Organoids. A. Experimental scheme for the infection of kidney organoids exposed to Control or Diabetic conditions. B. Immunofluorescence of Control and Diabetic kidney organoids at 1 day post infection (1 dpi) with Mock or SARS-CoV-2 (10^6^ virus particles/organoid as determined in Vero cells). Confocal images show the detection of ACE2 (green), virus nuclear protein (NP; red), LTL (magenta) and DAPI (blue). Scale bars, 250 μm, 50 μm (magnified views). *n = 2* organoids per condition. C. Quantification of SARS-CoV-2 infection in B) as % of NP occupied area. Data are shown as mean ± SEM. *n = 2* organoids per condition. D. mRNA expression levels of SARS-CoV-2 in Control and Diabetic kidney organoids at 1 dpi with Mock or SARS-CoV-2 (10^6^ virus particles/organoid as determined in Vero cells). See also Figure S3; Figure S4; Figure S5.

To assess potential transcriptional changes induced by SARS-CoV-2, we performed single cell RNA sequencing (scRNAseq) across 4 biological conditions (mock and 1dpi, in both 5 mM glucose- and high oscillatory glucose-treated kidney organoids). Cell types in kidney organoids were assigned using unsupervised clustering after integrating control versus diabetic infected conditions and the Uniform Manifold Approximation and Projection (UMAP) algorithm to visualize the scRNAseq data (Figure 4A). We retrieved renal endothelial-like, mesenchymal, proliferating, podocyte and tubule cell populations in all four experimental samples indicating that apparently neither diabetic conditions nor SARS-CoV-2 infections altered cell compositions in the kidney organoids (Figure S4A). Organoids from the all four conditions contained cells representative of a developing nephron including ENG^+^ and PECAM1^+^ endothelial-like cells, NPHS1^+^ and NPHS2^+^ podocytes, LRP2^+^ and SLC3A1^+^ proximal tubule cells, and CDH1^+^ distal tubule cells (Figure S4B), resembling second trimester human fetal kidney cell populations (Figure S4C). Diabetic kidney organoids at 1dpi again showed increased numbers of cells containing viral RNA compared to control organoids (Figure 4B), further confirming our results. As expected from an active infection, SARS-CoV-2 infections resulted in an increase of inflammatory-related processes in kidney organoids under control and diabetic conditions (Figure 4C). SARS-CoV-2 infections were also associated with downregulation of glycolysis-related processes in diabetic organoids (Figure 4C). In addition to altered glycolysis, differential gene expression analysis showed a strong correlation with inflammation (e.g. CXCL family genes) and diabetic-related pathways (e.g. CEBPD, STAT3) in the SARS-CoV-2 infected diabetic kidney organoids (Figure 4D; Figure S4D-F).

**Figure 4.**
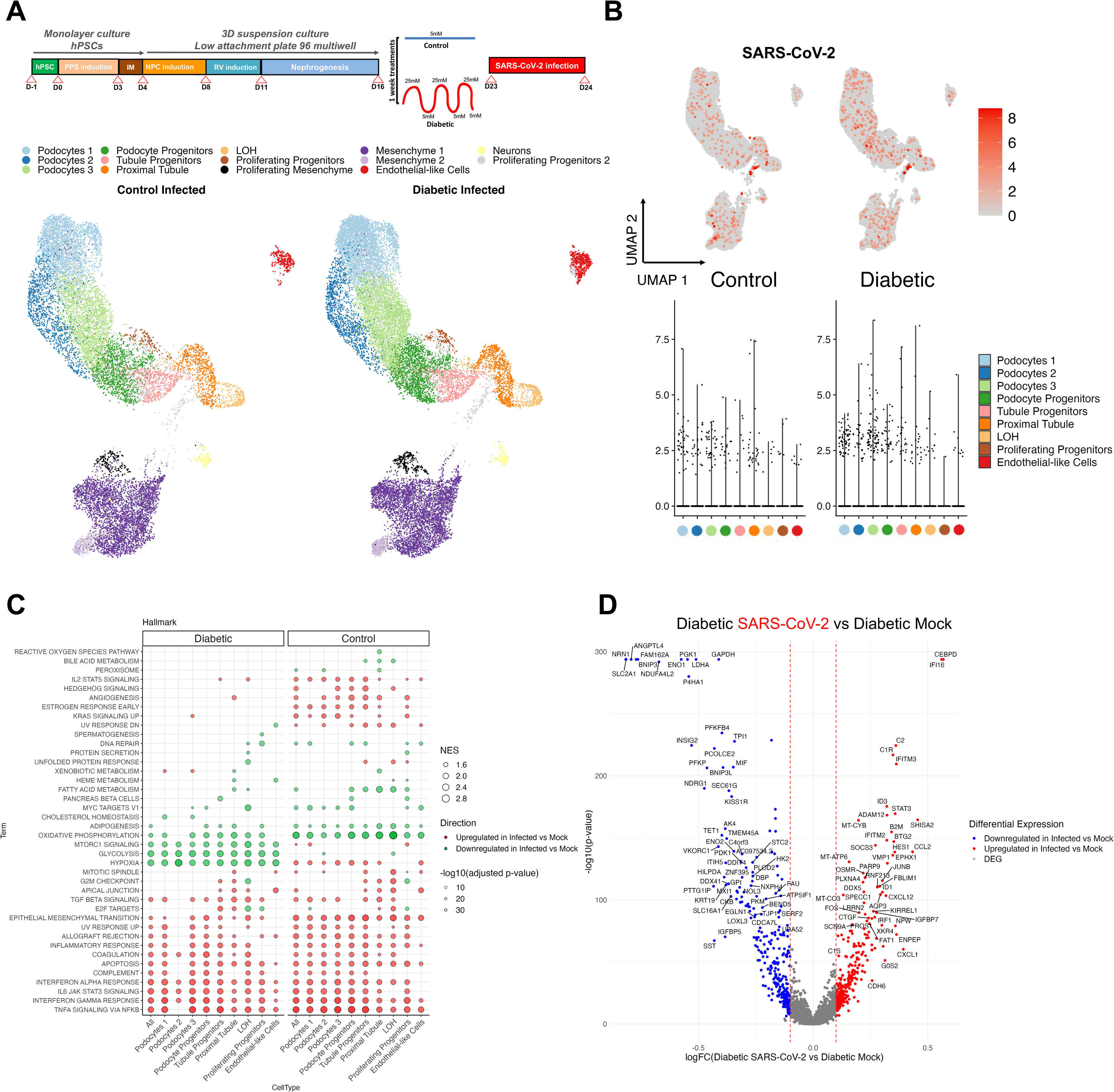
Metabolic Programing in Diabetic Human Kidney Organoids Boosts SARS-CoV-2 Infection. A. Schematics for the infection of kidney organoids exposed to Control or Diabetic conditions. Uniform manifold approximation and projection (UMAP) of Control and Diabetic kidney organoids at 1dpi with SARS-CoV-2 (10^6^ virus particles/organoid as determined in Vero cells). Clusters are colored by annotated cell types. B. UMAPs for SARS-CoV-2 expression in Control and Diabetic kidney organoids at 1dpi. Cells are colored based on expression level. The violin plots in the bottom panels represent expression level for the different cell types indicated in the legend. For SARS-CoV-2, expression is considered as zero for cells expressing < 5 UMIs. C. A hallmark GSEA was performed separately for Control and Diabetic conditions, comparing SARS-CoV-2 infected (10^6^ virus particles/organoid as determined in Vero cells) versus Mock organoids. The ten gene sets per direction and sample with lowest adjusted p-value are shown. Each column corresponds to one of the comparisons. Circles are coded by color (direction), size (NES) and transparency (p-value). D. Differentially expressed genes (DEGs) in the comparison of the SARS-CoV-2 infected (10^6^ virus particles/organoid as determined in Vero cells) against Mock organoids in Diabetic conditions considering only renal-like cell types. In the volcano plot, the x-axis indicates log fold change (FC) and the y-axis indicates statistical significance with the – log10(p-value). Genes with an adjusted p-value < 0.05 are considered upregulated (red) if the logFC > 0.1 and downregulated (blue) if the logFC < −0.1. Non DEG are shown in grey. See also Figure S4; Figure S5.

Recent findings showed that glycolysis sustains SARS-CoV-2 replication in human monocytes exposed to high glucose (11 mM) [29]. Considering our findings that SARS-CoV-2 infection in diabetic kidney organoids led to a decrease in glycolysis, we also investigated the impact of SARS-CoV-2 infection in kidney organoids exposed to 11 mM glucose culture conditions. SARS-CoV-2 infections did not alter cell clustering nor cell type proportions (Figure S5A,B) across mock and 1dpi kidney organoids grown in 11 mM glucose. Moreover, cells presenting high expression of viral RNA were mainly located in the loop of Henle and proximal tubule cell clusters, again co-localized to ACE2 expression (Figure S5C). Importantly, following SARS-CoV-2 infections, we observed a switch from OXPHOS to a glycolytic-based metabolism in the diabetic kidney organoids in agreement with recent findings in human monocytes (Figure S5D) [29]. Analysis for differentially expressed genes confirmed those findings together with enrichment in IFN- and TNF-, IL2/6 and mTORC1 signaling (Figure S5D-F).

### ACE2 is essential for SARS-CoV-2 infections in kidney organoids

The ACE2 receptor is sufficient for SARS-CoV-2 entry into human cells, though other receptors have been proposed [30]. Whether ACE2 is essential for SARS-CoV-2 infections is still not known. We and others have previously shown that ACE2 expression protects the lung from injury and that its expression is downregulated *in vivo* and *in vitro* upon SARS-CoV and SARS-CoV-2 spike protein engagement on the cell membrane [31–34]. Thus, we assessed ACE2 expression by qPCR in kidney organoids after SARS-CoV-2 infection. ACE2 mRNA levels were significantly downregulated after SARS-CoV-2 infection compared to mock conditions in kidney organoids in both control (5 mM glucose) and diabetic (high oscillatory glucose) regimes (Figure 5A). Of note, we found that at 1dpi cells expressing ACE2 as well as TMPRSS2, a cellular protease that cleaves and activates the SARS-CoV-2 envelope spike protein [35], were primarily located in the proximal tubule and loop of Henle (Figure S6A), respectively, as observed in the native kidney [27, 28].

**Figure 5.**
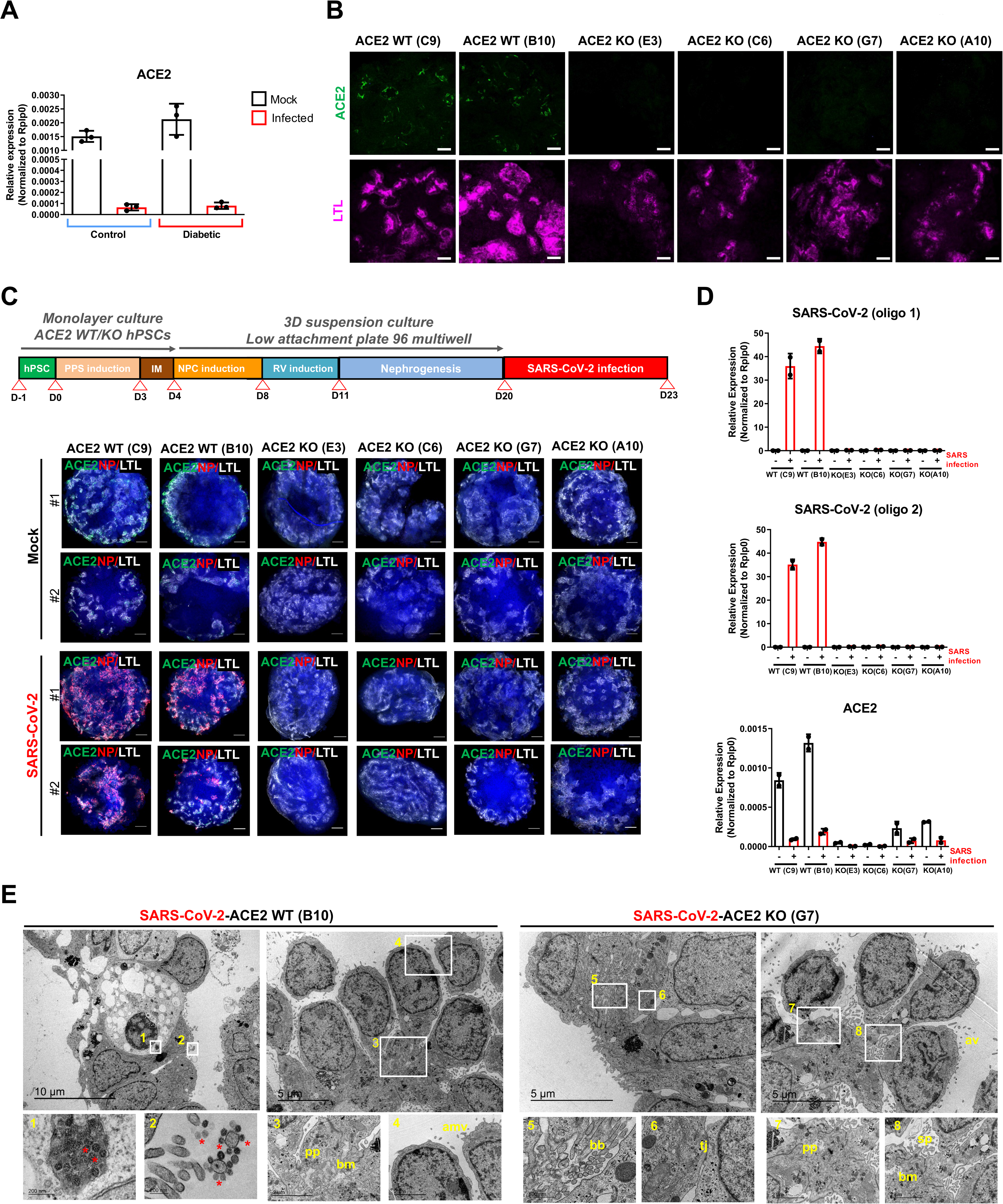
SARS-CoV-2 Infection in Human Kidney Organoids Depends on ACE2. A. mRNA expression levels of ACE2 in Mock or SARS-CoV-2 infected (10^6^ virus particles/organoid as determined in Vero cells) kidney organoids exposed to Control or Diabetic conditions. B. Immunofluorescence staining for ACE2 (green), and LTL (magenta) in ACE2 WT and ACE2 KO kidney organoids. Scale bars, 50 μm. C. Experimental scheme for the generation and viral infection of ACE2 WT and ACE2 KO kidney organoids. Immunofluorescence was performed at 3 days post infection (3 dpi) in Mock or SARS-CoV-2 infected (10^6^ virus particles/organoid as determined in Vero cells) specimens for the detection of ACE2 (green), virus nuclear protein (NP; red), LTL (grey) and DAPI (blue). Scale bars, 250 μm. D. Quantitative real time for the detection of SARS-CoV-2 and ACE2 mRNA was performed at 3 dpi in Mock or SARS-CoV-2 infected (10^6^ virus particles/organoid as determined in Vero cells) ACE2 WT and ACE2 KO kidney organoids. Data from a pool of 12 organoids/group is shown. E. TEM analysis of ACE2 WT and ACE2 KO kidney organoids infected with SARS-CoV-2 (10^6^ virus particles/organoid as determined in Vero cells) and recovered at 3 dpi. Representative images of infected ACE2 WT specimen show numerous viral particles (asterisks) inside a vesicle near the plasma membrane of a dying cell (1) and in the apical microvilli (amv) of a tubular-like cell (2). Details for podocyte-like cells exhibiting podocyte related-structures including primary processes (pp) (3), the deposition of a basement membrane (bm) (3), and apical microvilli (amv) (4) are shown. Scale bars, 10 μm and 5 μm; 200 nm (magnified views in 1 and 2); 2 μm (magnified views in 3 and 4). Representative images of infected ACE2 KO specimen show epithelial tubular-like cells with brush borders (bb) (5) and tight junctions (tj) (6). Details for podocyte-like cells with podocyte related-structures including primary (pp), the deposition of a basement membrane (bm), and cell processes (sp) are shown. Scale bars, 5 μm; 500 nm (magnified views in 5 and 6); 2 μm (magnified views in 7 and 8). See also Figure S6; Figure S7; Figure S8; Figure S9.

Our data showed that ACE2 is being upregulated at oscillatory high glucose exposure, but markedly downregulated following SARS-CoV-2 infections in kidney organoids. To test whether ACE2 is indeed essential for SARS-CoV-2 infections in kidney organoids, we generated ACE2 knockout (KO) hPSC lines using CRISPR/Cas9 genome editing (Figure S6B,C). Upon differentiation, H&E staining showed that both wild type (WT) and ACE2 KO kidney organoids exhibited similar nephron-like structures containing podocyte-like and tubule-like cells (Figure S6D). ACE2 deficiency was confirmed by immunofluorescence (Figure 5B), western blotting (Figure S6E), and qPCR (Figure S6F). scRNAseq profiling and qPCR analysis of WT and ACE2 KO kidney organoids showed no changes in renal cell populations, further confirming that the knockout of the ACE2 in hPSCs does not interfere with the renal differentiation process (Figure S6F and Figure S7A,B). Expression of TMPRSS2, was not changed in ACE2 mutant kidney organoids (Figure S6F). In addition, we found that expression of basigin (BSG, also known as CD147), proposed as another putative receptor for SARS-CoV-2 [36], was not affected by ACE2 deficiency (Figure S7C). Interestingly, ACE2 KO kidney organoids exhibited an upregulation in oxidative phosphorylation (OXPHOS) processes compared to ACE2 WT organoids as shown by gene set enrichment analysis (GSEA) (Figure S7D). These results are in line with previous observations that genetic deletion of ACE2 or pharmacological inhibition of ACE2 induces renal oxidative stress and promotes diabetic renal injury [37, 38].

Control WT and ACE2 KO kidney organoids were infected with SARS-CoV-2 and virus infection was monitored at 3 dpi (Figure 5C). Immunofluorescence analysis showed a robust SARS-CoV-2 infection in WT ACE2 organoids based in NP detection in LTL^+^ tubular-like cells (Figure 5C, Figure S7E). Infection was confirmed by the detection of SARS-CoV-2 mRNA by qPCR (Figure 5D). Immunofluorescence analysis showed that SARS-CoV-2 infection led to an increase in the expression of the kidney injury molecule 1 (KIM1) in both WT and ACE2 KO organoids (Figure S7E). Loss of ACE2 expression upon infection was re-confirmed at the RNA (Figure 5D) and protein level (Figure S7F). No changes in the levels of expression of podocyte marker genes (WT1, PODXL, NPHS1, NPHS2 and MAFB), nor for tubular marker genes (SLC3A1 and SLC16A1) were found after infection comparing WT and ACE2 KO kidney organoids (Figure S8A). TEM analysis confirmed that WT and ACE KO kidney organoids exhibited tubular-like cells and podocyte-like cells with the detection of SARS-CoV-2 viral particles in tubular-like cells in the WT background, corresponding to ACE2 expression (Figure 5E). Most importantly, ACE2 deletion resulted in a complete absence of NP expression in kidney organoids and almost undetectable levels of SARS-CoV-2 mRNA (Figure 5C,D; Figure S7E). TEM analysis confirmed the absence of viral particles in ACE2 KO kidney organoids (Figure 5E). These data provide the first genetic proof that ACE2 is the essential receptor for SARS-CoV-2 in kidney organoids.

### Inactivation of BSG/CD147 in kidney organoids has no effect on SARS-CoV-2 infections

In contrast to ACE2, SARS-CoV-2 infections did not lead to apparent changes in the levels of BSG/CD147 (Figure S7C), nor TMPRSS2 and NPR-1, another proposed co-factor involved in ACE2-mediated SARS-CoV-2 infection [39, 40] in WT or ACE2 KO kidney organoids, as determined by qPCR analysis (Figure S8A,B). To further validate the unique role of ACE2 in SARS-CoV-2 infection in kidney organoids we generated BSG KO hPSCs using CRISPR/Cas9 (Figure S9A,B). Upon differentiation, H&E staining showed that both the respective WT and BSG KO kidney organoids exhibited the presence of nephron-like structures containing podocyte-like cells and tubular structures (Figure S9C). BSG deletion was confirmed in BSG KO kidney organoids using qPCR (Figure S9D). To determine if BSG also plays a role in SARS-CoV-2 infection, as previously proposed, WT and BSG KO kidney organoids were infected with SARS-CoV-2 and analyzed at 3 dpi (Figure S9E). TEM analysis revealed the presence of viral particles in both WT and BSG KO organoids (Figure S9F). This was confirmed by qPCR analysis showing SARS-CoV-2 mRNA expression in both BSG KO and WT kidney organoids (Figure S9G) and immunofluorescence analysis to detect the viral NP (Figure S9H). These results do not support a role for BSG/CD147 in SARS-CoV-2 infections of human kidney organoids.

### ACE2 is essential for SARS-CoV-2 infections in diabetic kidney organoids

We next assessed whether ACE2 is essential for the observed increase in SARS-CoV-2 infections under our diabetic culture conditions. To address this question this, WT and ACE2 KO kidney organoids were exposed to 5 mM glucose (control) and high oscillatory glucose (diabetic) conditions and analyzed at 1 dpi (Figure 6A). Importantly, infected ACE2 KO kidney organoids were negative for viral NP irrespective of the control or diabetic treatment, contrary to ACE2 WT kidney organoids which again displayed significantly enhanced SARS-CoV-2 NP within the LTL^+^ proximal tubule structures when the kidney organoids were cultured under high oscillatory glucose conditions (Figure 6B). This was confirmed by qPCR analysis for the detection of SARS-CoV-2 mRNA (Figure 6C) and electron microscopy to determine viral particles (Figure S10). As expected, ACE2 mRNA and protein levels remained undetectable in the diabetic ACE2 KO kidney organoids (Figure 6D). qPCR analysis for the SARS-CoV-2 viral entry-associated genes TMPRSS2, BSG and NRP-1, and the renal markers NPHS1, PODXL and SLC3A1 showed no apparent changes irrespective of the control or diabetic treatment or ACE2 genetic background (Figure 6E). Thus, diabetic conditions license enhanced SARS-CoV-2 infections in kidney organoids, which is critically dependent on ACE2 expression.

**Figure 6.**
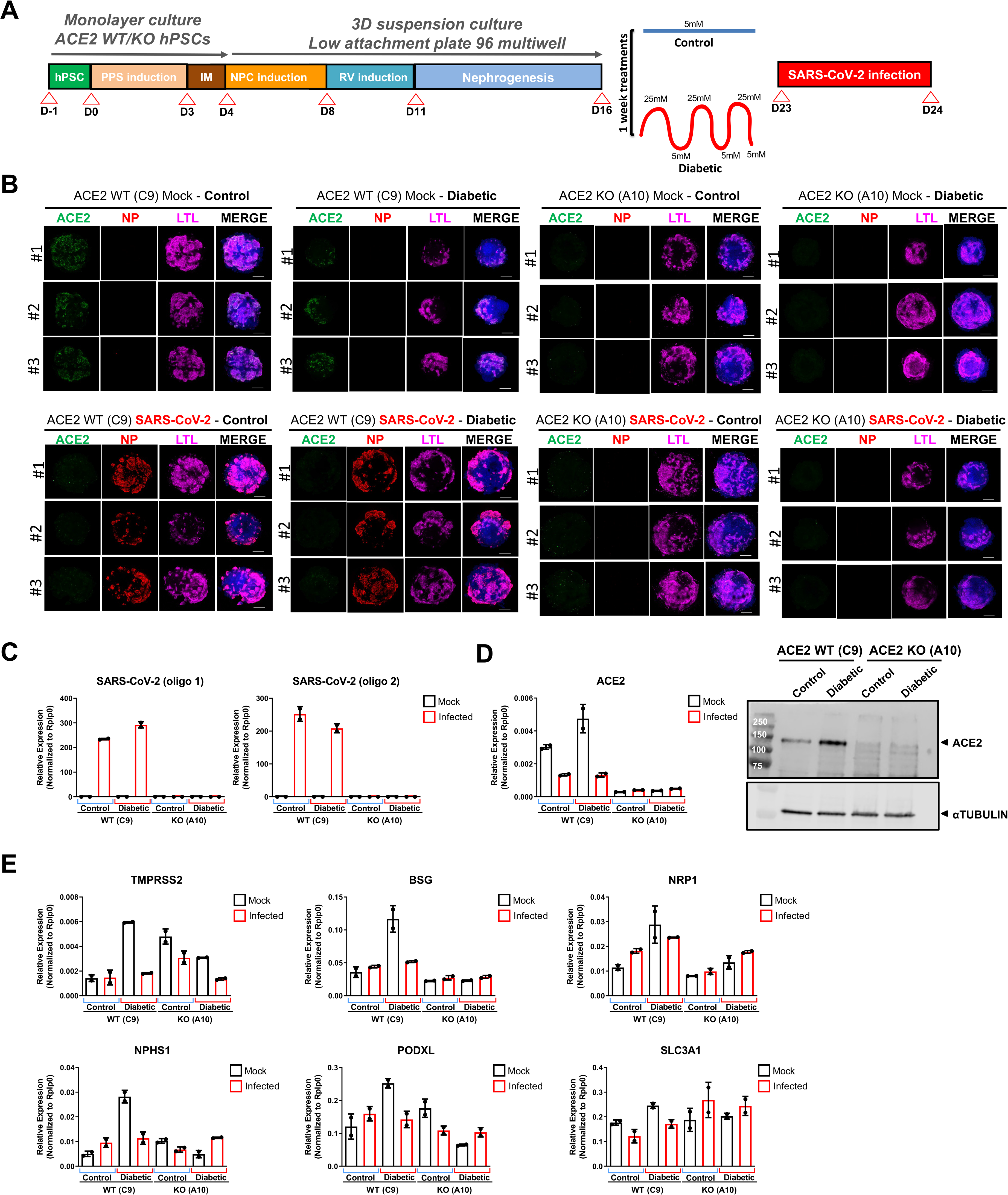
ACE2 Expression in Diabetic Human Kidney Organoids License Enhanced SARS-CoV-2 Infections. A. Experimental scheme for the infection of ACE2 WT and ACE2 KO kidney organoids exposed to Control or Diabetic conditions. B. Immunofluorescence was performed at 1 day post infection (1 dpi) in Mock or SARS-CoV-2 infected (10^6^ virus particles/organoid as determined in Vero cells) specimens exposed to Control or Diabetic conditions for the detection of ACE2 (green), virus nuclear protein (NP; red), LTL (magenta) and DAPI (blue). Scale bars, 250 μm. *n = 3* organoids per condition. C. Quantitative real time for the detection of SARS-CoV-2 mRNA was performed at 1 dpi in Mock or SARS-CoV-2 infected (10^6^ virus particles/organoid as determined in Vero cells) ACE2 WT and ACE2 KO kidney organoids exposed to Control or Diabetic conditions. Data from a pool of 12 organoids/group is shown. D. Quantitative real time for the detection of ACE2 mRNA was performed at 1 dpi in mock or SARS-CoV-2 infected (10^6^ virus particles/organoid as determined in Vero cells) ACE2 WT and ACE2 KO kidney organoids exposed to Control or Diabetic conditions (left). Data from a pool of 12 organoids/group is shown. Protein levels of ACE2 in Mock and SARS-CoV-2 infected (10^6^ virus particles/organoid as determined in Vero cells) ACE2 WT and ACE2 KO kidney organoids under Control or Diabetic conditions by Western Blot analysis (right). aTUBULIN was used as loading control. Data from a pool of 12 organoids/group is shown. E. Quantitative real time was performed at 1 dpi in Mock or SARS-CoV-2 infected (10^6^ virus particles/organoid as determined in Vero cells) ACE2 WT and ACE2 KO kidney organoids under Control or Diabetic conditions for the indicated markers. Data from a pool of 12 organoids/group is shown. See also Figure S10.

### Enhanced SARS-CoV-2 infection in kidney cells from diabetic patients

It is still unclear whether SARS-CoV-2 can directly infect renal cells in the human kidney or if the damage observed in this organ are secondary due to COVID-19 [41–44]. Although our data unequivocally show that kidney organoids can be infected via ACE2, we wanted to also determine SARS-CoV-2 infections of primary human renal cells. We therefore isolated kidney human proximal tubular cells (HPTC) from kidney biopsies of non-diabetic (control) and diabetic patients (Figure 7A). The mRNA levels of PGC1α were significantly downregulated in diabetic HPTCs compared to control counterparts (Figure 7B), paralleling our findings in the diabetic organoids and diabetic kidney organoids-isolated LTL^+^ cells (Figure S1G; Figure S2C). In addition, HPTCs isolated from diabetic kidney biopsies exhibited higher OCR, basal respiration, ATP production and maximal respiratory capacity compared to control counterparts (Figure 7C). Importantly, following SARS-CoV-2 infections, we detected increased numbers of NP^+^ cells in diabetic HPTCs as compared to renal cells isolated from kidneys of non-diabetic controls (Figure 7D,E). Moreover, diabetic patient samples had markedly higher amounts of viral RNA as detected by qPCR (Figure 7F) and produced a larger amount of infectious viral particles as determined by the Median Tissue Culture Infectious Dose (TCID50) assay (Figure 7G). Of note, we also observed increased ACE2 expression on HPTCs from diabetic patients (Figure 7H). Thus, human kidney proximal tubular cells can be directly infected with SARS-CoV-2 and diabetes licenses kidney cells to increased infections.

**Figure 7.**
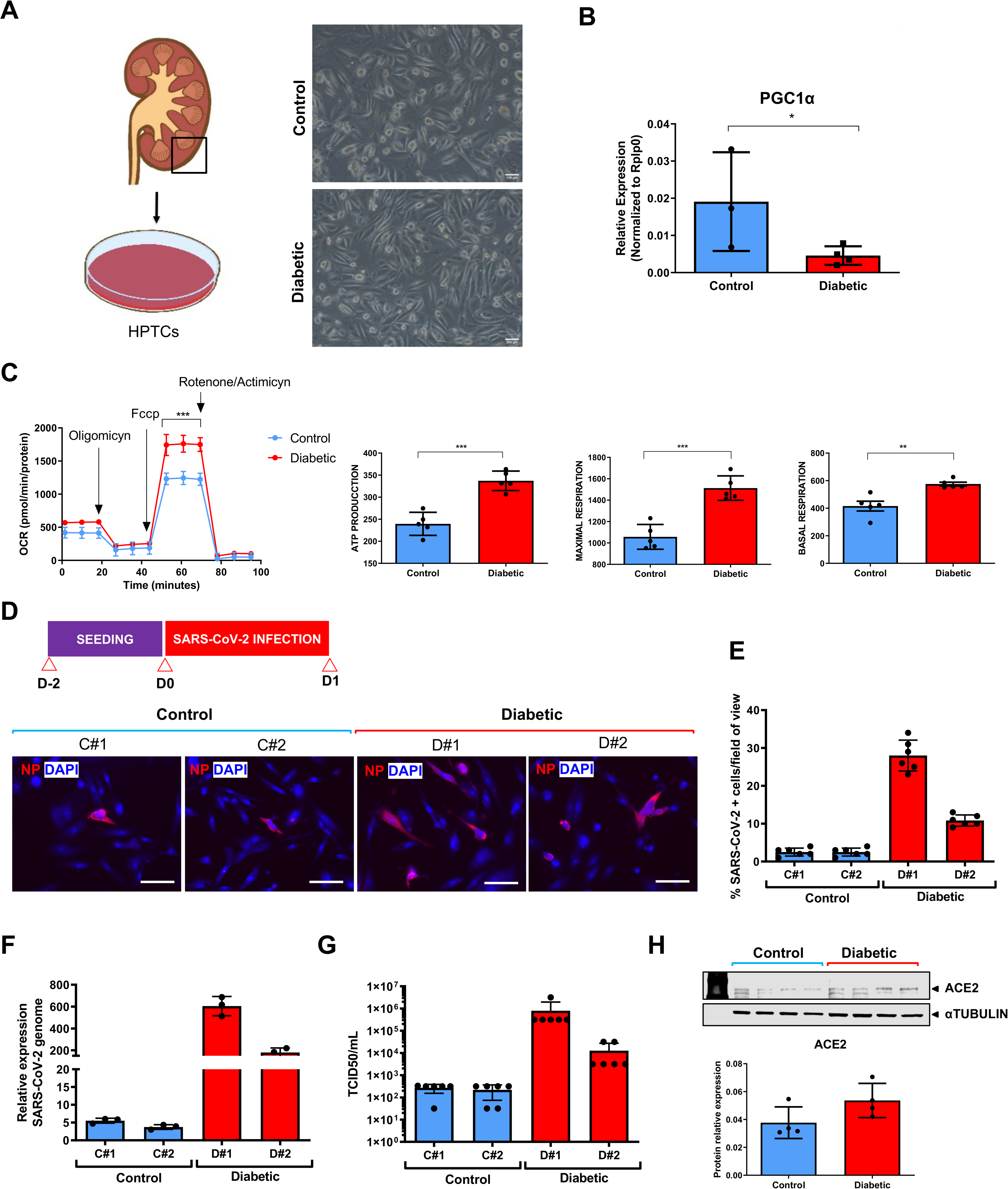
SARS-CoV-2 Infection in Tubular Epithelial Cells Derived from Diabetic Human Kidney Biopsies. A. Representative bright field images of human proximal tubular epithelial cells (HPTCs) isolated from kidney biopsies from non-diabetic (Control) and diabetic patients (Diabetic). Scale bars, 100 μm. B. mRNA expression levels of PGC1a by qPCR in Control and Diabetic HPTCs. Mean ±SEM is calculated from 3 (Control) and 4 (Diabetic) biological replicates. C. Kinetic oxygen consumption rate (OCR) response, basal respiration and spare respiratory capacity, cellular ATP production and inner mitochondrial membrane proton leak of HPTCs from Control and Diabetic HPTCs. Data are normalized to total protein. Mean ± SEM is calculated from 1 biological replicates/group performing 5 technical replicates each. **p<0.01; ***p<0.001***, two-way ANOVA, followed by Bonferroni post-test. D. Experimental scheme for viral infection in Control and Diabetic HPTCs. Immunofluorescence in SARS-CoV-2 infected (MOI 1 as determined in Vero cells) Control and Diabetic HPTCs at 1 dpi for the detection of virus nuclear protein (NP, red). Scale bars, 100 μm. E. Quantification of changes in NP protein expression in D) is shown as percentage of SARS-CoV-2 positive cells (+) per field. F. qPCR was performed at 1 dpi in SARS-CoV-2 infected (MOI 1 as determined in Vero cells) Control and Diabetic HPTCs for the detection of SARS-CoV-2 mRNA. G. Tissue culture infectious dose (TCID50)/mL is shown at 1 dpi SARS-CoV-2 infected (MOI 1 as determined in Vero cells) Control and Diabetic HPTCs. H. Protein levels of ACE2 in HPTCs from Control and Diabetic donors are shown by Western Blot. aTUBULIN was used as loading control.

## Discussion

Diabetes has emerged as one of the most frequent co-morbidities associated with severity and mortality of COVID-19. Given the current COVID-19 pandemic, there is an urgent need to better understand the complex relationships between pre-existing conditions in COVID-19 patients which may exacerbate viral infection and disease outcomes. In this regard, why diabetic individuals are more prone to develop severe COVID-19 remained largely unknown. Here, we developed a unique and novel culture system to generate diabetic human kidney organoids, based on oscillation of glucose levels also observed in DM patients. Our approach preserved renal cell types while recapitulating early hallmarks of diabetic kidney disease, including changes in ECM composition and metabolic reprogramming [45]. Using our system we could directly demonstrate that diabetic conditions enhanced SARS-CoV-2 infections, which is critically dependent on ACE2 expression.

We analyzed kidney organoids for expression of the SARS-CoV-2 entry receptor ACE2 and consistently observed ACE2 positivity in tubular-like cells by confocal microscopy, confirming our previous observations using single cell profiling in human kidney organoids [10] and earlier reports using single cell profiling [46] and immunohistochemistry [47] in the mouse and human kidney [48]. Recently single cell profiling from 436 patients suggested that increases in ACE2 expression within lungs and kidneys may increase the risk of SARS-CoV-2 infections [49]. Our results now show that a high oscillatory glucose regimen induces the expression of ACE2 at both the mRNA and protein levels consistent with previous findings showing increased ACE2 levels in the kidney cortex from db/db mice and STZ diabetic mice [50] and proximal tubular cells in kidney biopsies from patients with diabetic kidney disease [51]. Importantly, concurrent with enhanced ACE2 expression we observed increased viral loads at the mRNA and protein levels in the diabetic kidney organoids. Following infection, we detected a marked decrease in ACE2 mRNA expression in accordance with previous results using colon- and ileum-derived human intestinal organoids [34]. Single cell profiling showed a decrease in OXPHOS and decrease in glycolytic-based metabolism as well as a hypoxic signature in response to SARS-CoV-2 infection in diabetic organoids. Moreover, we observed alterations in mTORC1. Interestingly, the mTORC1 pathway, a master regulator of anabolic metabolism, has been recently highlighted as a candidate for COVID-19 therapeutic intervention [52, 53].

Nearly two decades ago, we and others have shown in *ace2* mutant mice that ACE2 is a negative regulator of the RAS system and genetically controls cardiovascular function and damage of multiple organs such as the lung, liver, and kidney [54, 55]. From the beginning of the pandemic ACE2 took the center stage in the COVID-19 outbreak as a receptor for the spike glycoprotein of SARS-CoV-2 [21, 56] which spurred the development of vaccines and therapies targeting the ACE2-SARS-CoV-2 Spike interaction. Since various other candidate receptors have been reported (and considering the drug and vaccine development landscape) it is therefore paramount to establish whether ACE2 is not only sufficient for infection but is in fact essential. Surprisingly, this has never been shown using human tissue or in experimental animal studies. To evaluate the key role of ACE2 in SARS-CoV-2 infection we therefore generated ACE2 knockout hPSCs. ACE2 KO kidney organoids, derived from these mutant cells, showed preserved renal differentiation; though, in line with the previously reported functions of ACE2, these ACE2 mutant organoids displayed differences with regards to OXPHOS, lipid metabolism as well as angiogenesis/endothelium [57]. Importantly, deletion of ACE2 completely prevented SARS-CoV-2 infections in control as well as diabetic kidney organoids, providing first genetic evidence on the essential role of ACE2 in SARS-CoV-2 infections in an engineered human tissue. To further demonstrate the unique role of ACE2 for viral entry, we also generated BSG/CD147 KO hPSCs lines as others have highlighted its role in SARS-CoV-2 infection [36, 58]. However, BSG KO kidney organoids supported viral infection, excluding an important role of BSG in renal SARS-CoV-2 infections. Multiple other cell types express ACE2, including intestinal cells or cardiomyocytes and heart alterations were the first phenotype we observed in *ace2* mutant mice [55]. It will be important to expand our studies to other organoid systems and other reported candidate SARS-CoV-2 entry receptors such as Neuropilin-1 (NRP1), to assess whether ACE2 is also essential in these engineered tissues.

For many of the systemic manifestations of COVID-19 it is unclear whether the pathology is a secondary “side effect” of the SARS-CoV-2 infection, such as immune activation or altered coagulation, or whether it is also due to a direct SARS-CoV-2 infection of specific organs. We first described that kidney organoids can shed progeny SARS-CoV-2 viruses [10, 59] and later investigations confirmed SARS-CoV-2 kidney tropism [41, 42], including the ability to replicate in human kidney cells, thereby establishing an association of kidney infection by SARS-CoV-2 with shorter survival time and increased incidence of acute kidney injury in COVID-19 patients [41, 42]. However, it should be noted that some studies failed to detect SARS-CoV-2 in kidneys [43, 44]. Our results now indicate that imbalances in cellular metabolism and ACE2 expression in kidney organoids due to elevated glucose levels directly lead to higher viral loads upon infection potentially leading to a switch from an OXPHOS to an aerobic glycolytic state that could further contribute to the severity of disease. We also observed increased SARS-CoV-2 infections in kidney proximal tubular cells isolated from diabetic patients, confirming our data in diabetic kidney organoids. Our results provide an experimental blueprint to dissect the role and contribution of co-morbidities in SARS-CoV-2 infections and COVID-19 pathogenesis. In particular, we here establish a link between imbalances in cellular metabolism in diabetes with ACE2 expression and the severity of SARS-CoV-2 infections.

### Limitations of the Study

Future work will be needed to address limitations of our study. (1) Utilizing more complex culture systems, including the assembly of a vascular component to kidney organoids, would help to further explore the impact of a high oscillatory glucose regimen in a more physiologically relevant context. (2) The design of our studies focused on the early stages of infection. As such, we cannot make any predictions with respect to the observed changes at later stages of the disease process. In this regard, inflammation represents a complex network of pathways which are influenced by external processes which currently cannot be simulated in our model. (3) We did not study pancreatic organoids, and the pancreas is one of the major metabolic organs that could explain later complications of COVID-19.

## Acknowledgements

We thank all members from our laboratories for helpful discussions and technical support. This work has received funding from the European Research Council (ERC) under the European Union’s Horizon 2020 research and innovation Programme (StG-2014-640525_REGMAMKID to P.P., and N.M.). NM is also supported by the Spanish Ministry of Economy and Competitiveness/FEDER (SAF2017-89782-R), the Generalitat de Catalunya and CERCA Programme (2017 SGR 1306), Asociación Española contra el Cáncer (LABAE16006) and Institute of Health Carlos III (ACE2ORG and PTC20/00013). N.M., E.G. and C.H.P. are supported by EFSD/Boehringer Ingelheim European Research Programme in Microvascular Complications of Diabetes. This research has been supported by EIT Health under grant ID 20366 (R2U-Tox-Assay) to E.G. and N.M. C.H.P. is supported by Marie Skłodowska-Curie Individual Fellowships (IF) grant agreement no. 796590. This work was supported in part by the ISCIII and co-financed by FEDER through TERCEL RETIC RD16/0011/0005 and RD16/0011/0027, CIBERONC CB16/12/00489; Gobierno de Navarra Departamento de Desarrollo Economico y Empresarial AGATA (0011-1411-2020-000011), DIANA (0011-1411-2017-000029) to F.P., A.V-Z. and A.U-A. F.G. is supported by the Ramon y Cajal Grant -Biomedicine (RYC-2014-16751) from the Ministry of Economy and Competitivity (MINECO), Spain. A.M. is supported by the IBEC International PhD Programme “La Caixa” Severo Ochoa fellowships (LCF/BQ/SO16/52270019). This project has received funding from the European Union’s Horizon 2020 research and innovation programme under grant agreement no. 878827 BRAV3 to N.M. and C.T. J.M.P. and the research leading to these results has received funding from the T. von Zastrow foundation, the FWF Wittgenstein award (Z 271-B19), the Austrian Academy of Sciences and the Canada 150 Research Chairs Program F18-01336 as well as the Canadian Institutes of Health Research COVID-19 grants F20-02343 and F20-02015. A.M. is supported by Swedish research council (2018-05766). This project has received funding from the Innovative Medicines Initiative 2 Joint Undertaking (JU) under grant agreement no. 101005026. The JU receives support from the European Union’s Horizon 2020 research and innovation program and EFPIA (A.M., J.M.P. and N.M.). This research has been supported by the project COV20/00278 from Instituto de Salud Carlos III to A.M., A.U., F.P. J.M.P., and N.M. This project has received funding from Ayudas Fundación BBVA a Equipos de Investigación Científica SARS-CoV-2 y COVID-19 (A.M, J.M.P and N.M).

## Author Contributions

This study was conceived and designed by N.M. N.M. and J.M.P. wrote the paper. E.G., P.P., C.H.P. performed all cell culture experiments in wild type cell lines. E.G. and P.P. performed all cell culture experiments in genome edited cell lines. E.G. and P.P. performed all histological analysis. E.G. performed TEM analysis. E.G., P.P., C.H.P., G.J., A.L., A.H., R.O., D.M-R., C.T., P.D-P., O.H., J.M.P., and N.M., performed organoid data analysis including qPCR, western blot and image quantification. M.S., V.M., S.B. and A.M. performed all the experiments involving SARS-CoV-2, including isolation, and helped with manuscript editing. F.G. and A.M. performed CRISPR/Cas9 lines. M.S., S.B. and A.V-Z. prepared scRNAseq samples. A.U-A., J.P.R and F.P. performed all the single cell data analysis on genome edited derived organoids. E.G., D.M-R., P.A-V., J.M.C. and N.M. performed patient sample isolation and characterization.

## Declaration of Interests

A patent has been submitted to use human organoids to study SARS-CoV-2 infections and possibly develop new therapies. J.M.P. is shareholder of Apeiron Biologics that is developing soluble ACE2 for COVID-19 therapy.

## Notes

https://www.ncbi.nlm.nih.gov/geo/query/acc.cgi?acc=GSE181002

