## Supplementary material for "A diabetic *milieu* increases cellular susceptibility to SARS-CoV-2 infections in engineered human kidney organoids and diabetic patients"

[1] Pluripotency for Organ Regeneration, Institute for Bioengineering of Catalonia (IBEC), The Barcelona Institute of Science and Technology (BIST), Barcelona, Spain. [2] Department of Infectious Diseases, Molecular Virology, Heidelberg University Hospital, Heidelberg, Germany. [3] Research Group “Cellular Polarity and Viral Infection”, German Cancer Research Center (DKFZ), Heidelberg, Germany. [4] Karolinska Institute and Karolinska University Hospital, Unit of Clinical Microbiology, SE-17182, Stockholm, Sweden. [5] Área de Hemato-Oncología, Centro de Investigación Médica Aplicada, Instituto de Investigación Sanitaria de Navarra (IDISNA), Universidad de Navarra, 31008 Pamplona, Spain. [6] Centro de Investigación Biomédica en Red de Cáncer (CIBERONC), 28029 Madrid, Spain. [7] Departamento de Hematología, Clínica Universidad de Navarra, Universidad de Navarra, 31008 Pamplona, Spain. [8] IMBA, Institute of Molecular Biotechnology of the Austrian Academy of Sciences, Dr. Bohr-Gasse 3, 1030 Vienna, Austria. [9] Center for Bioengineering and Tissue Regeneration, UCSF, San Francisco, California, USA. [10] Internal Medicine Department, Hospital Universitario de la Santa Creu i Sant Pau, Barcelona, Spain. [11] Department of Medical Genetics, Life Sciences Institute, University of British Columbia, Vancouver, British Columbia, Canada. [12] Nephrology and Kidney Transplant Department, Hospital Clínic Barcelona, Barcelona, Spain. [13] Laboratori Experimental de Nefrologia I Trasplantament (LENIT), Fundació Clínic per a la Recerca Biomèdica (FCRB), Barcelona, Spain. [14] National Veterinary Institute, Uppsala, Sweden. [15] Department of Infectious Diseases, Virology, Heidelberg University Hospital, Heidelberg, Germany. [16] Catalan Institution for Research and Advanced Studies (ICREA), Barcelona, Spain. [17] Centro de Investigación Biomédica en Red en Bioingeniería, Biomateriales y Nanomedicina, Madrid, Spain. [18] Lead Contact.

‡ these authors equally contributed to the development of the work presented here.

***Corresponding authors:**

Nuria Montserrat

Pluripotency for Organ Regeneration Laboratory

Institute for Bioengineering of Catalonia (IBEC), the Barcelona Institute of Science and Technology (BIST)

08028 Barcelona, Spain

Josef Penninger

Department of Medical Genetics, Life Sciences Institute, University of British Columbia

Vancouver, British Columbia, Canada

Steeve Boulant

Department of Infectious Diseases, Virology, Heidelberg University Hospital,

Heidelberg, Germany

Ali Mirazimi

Karolinska Institute and Karolinska University Hospital, Unit of Clinical Microbiology,

SE-17182, Stockholm, Sweden

#### Material and Methods

##### *Cells*

hPSC lines were obtained after the approval of the Ethics Committee from the Clinical Translational Program for Regenerative Medicine in Catalonia (P-CMR [C])and the Comisión de Seguimiento y Control de la Donación de Células y Tejidos Humanos del Instituto de Salud Carlos III (project numbers: 0336E/1517/2015; 0336E/4166/2020; 0336E/4168/2020; 0336E/1124/2021/ 0336E/2723/2021). hESC ES[4] line and CBiPS1sv-4F-40 iPSC line were obtained from The National Bank of Stem Cells (ISCIII, Madrid). Both hPSC lines were maintained and grown in Essential 8 medium (A1517001, ThermoFisher) in cell culture plates coated with 5 µg/mL vitronectin (A14700, ThermoFisher) with 5% CO2 at 37 °C. Cells were passaged every 4–6 days by disaggregating hPSC colonies into small cell clusters using 0.5 mM EDTA (15575-038, Thermofisher). Vero E6 (ATCC CRL 1586) were maintained in Dulbecco's modified Eagle's medium (DMEM) (Gibco) supplemented with 10% fetal bovine serum (FBS; 10270-106, Gibco) and 1% penicillin/streptomycin (15140122, ThermoFisher). Primary renal proximal tubular epithelial cells were isolated as previously described [60, 61] upon ethics committee of "Hospital Clinic de Barcelona" approved the procedure (project number: HCB/2021/0119). Primary renal proximal epithelial cells were plated on plastic plates in proximal tubular cell media consisting of DMEM without glucose (11966025, Thermofisher) supplemented with 5% FBS (Gibco), 5 mM glucose (Merck), penicillin/streptomycin (15140122, ThermoFisher), ITS 1X (I3146, Merck), and 40 ng/mL human EGF (236-EG-01M, R&D systems).

##### *SARS-CoV-2 isolate*

SARS-CoV-2 were isolated on Vero-E6 cells from nasopharyngeal sample of patient diagnosed with COVID-19 in Sweden as described before in Monteil et al., 2020 [10]. Alternatively, SARS-CoV-2 (strain BavPat1) was obtained via the European Virology Archive. Virus were amplified in Vero E6 cells and titered using a plaque assay as previously described [62] with fixation of cells 72 hours post infection. Virus was used at a passage 3.

##### *Generation of genome edited hPSC lines*

*ACE2* and *BSG* knockout lines were generated using engineered ES[4] and CB[40] PSC lines, in which a doxycycline-inducible Cas9 cassette was introduced at the AAVS1 safe harbor locus [63, 64]. Two crRNAs per gene were designed using the Alt-R Custom Cas9 crRNA

Design Tool (IDT) (Key Resources Table). IDT crRNA/tracrRNA duplexes were annealed following manufacturer's guidelines and transfected using Lipofectamine RNAiMAX Transfection Reagent (13778150, Thermofisher) at 10 nM final concentration in 0.25 million doxycycline-treated ES[4] or CB[40] PSCs/mL. Cells were grown 3-5 days after transfection and single-cell seeded at 10, 20, 50, 100 cells/cm$^2$. Leftover cells were used for DNA purification and further pool editing analysis. Clonal lines were established by manual colony picking (48 clones/crRNA) and further replica-plated. One replica was stored in nitrogen, while the other was used for DNA purification and MiSeq analysis of a locus-specific pooled PCR library generated through two rounds of PCR introducing Illumina adapters and specific indexes labeling each clone (Key Resources Table). Indel frequency in edited pools and allelic composition of clones was obtained by analyzing trimmed Fastq reads using CRIPRESSO2 webtool [65]. For each crRNA edit, at least one non-edited clone (+/+), one heterozygous clone (+/frameshift, or +/fs), and one homozygous or trans-heterozygous clone (fs/fs) were amplified, stocked, and used for further experiments.

##### ***Kidney organoid differentiation***

Prior differentiation (day -1), undifferentiated hPSCs colonies were dissociated into cell clumps using 0.5 mM EDTA (15575-038Thermofisher) at 37°C for 3 minutes. To obtain single cell suspensions samples were then incubated in AccumaxTM (07921, Stem Cell Technologies) at 37°C for 3 additional minutes. Cells were counted using the Countess ® Automated Cell Counter (Invitrogen) and seeded at 100,000 cells/well on 24 multi-well plates coated with 5 µg/mL vitronectin in the presence of Essential 8 medium (A1517001, Life Technologies) at 37°C overnight. The next day (day 0), the differentiation was initiated by treating monolayer cultures with 8 µM CHIR99021 (CHIR; SML1046, Merck) in Advanced RPMI medium consisting of Advanced RPMI 1640 basal medium (12633020, ThermoFisher) supplemented with 1% Penicillin-Streptomycin and 1% of GlutaMAX$^{TM}$ (35050061, ThermoFisher) for 3 days with daily medium changes. On day 3, monolayer cultures were treated with 200 ng/mL FGF9 (100-23, Peprotech), 1 µg/mL heparin (H3149, Merck) and 10 ng/mL activin A (Act A) (338-AC-050, Vitro) in Advanced RPMI 1640 basal medium supplemented with 1% Penicillin-Streptomycin and 1% of GlutaMAX$^{TM}$ for 1 day. On day 4, cell monolayers were treated with 5 µM CHIR, 200ng/mL FGF9 and 1 µg/mL Heparin Advanced RPMI 1640 basal medium (12633020, ThermoFisher) supplemented with 1% Penicillin-Streptomycin and 1% of of GlutaMAX$^{TM}$ for 1 hour. After this 1 hour-treatment cell monolayers were dissociated using TrypLE™ Express Enzyme (1260402, ThermoFisher) for 1 min collected, and counted. Next, 100,000 cells/well were dispensed on a V-shape 96 multi-well plate (249935, ThermoFisher) and centrifuged at 300g for 5min. Cell spheroids were cultured in Advanced RPMI 1640 basal medium supplemented with 1% Penicillin-Streptomycin and 1% of GlutaMAX™, 200ng/mL FGF9 and 1 µg/mL heparin for 7 days with medium changes every other day. From day 11, organoids were maintained in Advanced RPMI 1640 basal medium supplemented with 1% Penicillin-Streptomycin and 1% of GlutaMAX™. From that stage medium was changed every other day.

***Glucose challenge in kidney organoids***

Prior treatments, day 16 kidney organoids were incubated with DMEM glucose-free (11966025, ThermoFisher) supplemented with 1% Penicillin-Streptomycin for 6h (starvation period). Normoglycemic conditions were emulated exposing kidney organoids to DMEM glucose-free (11966025, ThermoFisher) supplemented with 5 mM D-Glucose (G7021, Merck) changing medium every day until day 23. High oscillatory glucose conditions involved the exposure of day 16 kidney organoids to DMEM glucose-free (11966025, ThermoFisher) supplemented with 5mM D-Glucose (5 mM glucose medium) on even days or with 25mM of D-Glucose (25 mM glucose medium) on odd days until day 23 (5-25mM oscillatory glucose regime). 5 mM glucose medium was additionally supplemented with 20 mM D-Mannitol (M9647, Merck) to obtain the same molarity as in the 25 mM glucose medium.

***Isolation of proximal tubular cells from kidney organoids***

Kidney organoids were stained with fluorescein-conjugated LTL (FL-1321, Vector Laboratories) as described elsewhere [66]. Kidney organoids were then dissociated to single cells using Accumax (07921, Stem Cell Technologies) for 15 min followed by 0.25% (wt/vol) trypsin (25300–054, Life Technologies) for 15 min at 37 °C. SA3800 software version 2.0.4 (SONY) was used to acquire flow cytometry samples in the Sony SA3800 spectral cell analyser (SONY). FACSDiva software version 8.0.1 (BD Biosciences) was used in the FACS Aria Fusion instrument (BD Biosciences) for cell sorting experiments. FlowJo software version 10 was used to analyse the data.

***ACE2 mRNA half-life in 5mM and 25mM cultured kidney organoids***

Kidney organoids were treated with actinomycin D to inhibit transcription. At the indicated times (0, 2 h, 4 h, and 8 hours) total RNA was isolated from the kidney organoids (two pools of 8 kidney organoids per time point) and relative ACE2 mRNA levels were determined by RT-qPCR as describe.

***SARS-CoV-2 infections of kidney organoids and proximal tubular kidney cells***

Kidney organoids were infected with $10^6$ SARS-CoV-2 infectious particles (as determined in Vero cells) in Advanced RPMI medium (Thermofisher) in a volume of 50µl per well of a 96-

well ultra-low attachment plate for 1 hour. Proximal tubular cells were seeded in 24-well plates in proximal tubular cell medium. One day post-seeding, cells were infected with $10^6$ MOI 1 (as determined in Vero cells) SARS-CoV-2 infectious particles in a final volume of 200µl per well in proximal tubular cell medium at 37°C for 1 hour followed by washing with PBS and adding 500µl of new proximal tubular cell medium. At the different days post-infection, samples (organoids or cells) were washed 3 times with PBS and then lysed using Trizol™ (Thermofisher), RIPA buffer (Thermofisher) or 4% paraformaldehyde (153799, Aname) for ulterior analysis.

***Single-cell RNA sequencing***

Single cell RNA sequencing was performed as described in our previous study [10]. Briefly, kidney organoids were collected and washed twice with PBS, and further dissociated into a single cell suspension by treating them with Accumax (07921, Stem Cell Technologies) for 15 min at 37ºC followed by Trypsin-EDTA 0.25% (wt/vol) trypsin (25300-054, Life Technologies) for additional 15 min at 37ºC. The reaction was deactivated by adding 10% FBS. The cell suspension was then passed through a 40µm cell strainer. After centrifugation at 1,000 RPM for 5 minutes cell numbers and viability were analyzed using Countess Automated Cell Counter (Invitrogen). Then cell suspensions were loaded onto a well of a 10x Chromium Single Cell instrument (10x Genomics). Barcoding and cDNA synthesis were performed according to the manufacturer's instructions. Qualitative analysis was performed using the Agilent Bioanalyzer High Sensitivity assay. The cDNA libraries were constructed using the 10x Chromium™ Single cell 3' Library Kit v3.1 according to the manufacturer's original protocol. Libraries were sequenced on either Illumina NovaSeq 6000 or NextSeq 500 500 2x150 paired-end kits using the following read length: 28bp Read1 for cell barcode and UMI, 8bp I7 index for sample index and either 94bp Read2 in the 5 mM, 11 mM and 5-25 mM oscillatory glucose samples (diabetic samples). For ACE2 WT and ACE 2KO samples (ACE2 samples) 55bp read length was used for Read2. Libraries were pre-processed using Cell Ranger (4.0.0) from 10X Genomics (http://10xgenomics.com). Reads from the ACE2 samples were aligned to the reference human genome (GRch38), whilst diabetic samples were aligned to a custom reference built for this work including the SARS-CoV-2 genome (isolate Wuhan-Hu-1, NC_045512.2). Genome annotation corresponded to Ensembl v93, adding a single gene with a single exon spanning the entire SARS-CoV-2 sequence. The median number of unique molecular identifiers (UMIs) per cell was between 2,300 and 4,600, with a median of 1,300-2,200 genes detected per condition.

##### ***Protein extraction and western blot analysis in kidney organoids***

Protein was extracted from kidney organoids cultured in 5 mM glucose, 5-25 mM oscillatory glucose or Advance RPMI with or w/o infection using RIPA buffer (ThermoFisher) supplemented with complete protease inhibitor cocktail (ThermoFisher). Samples were centrifuged at 13,000 g for 15 mins at 4°C. Supernatant were collected and protein concentration was measured using a bicinchoninic acid (BCA) protein quantification kit (Thermo Scientific). For western blot analyses, 25 ug of protein were separated in 10% sodium dodecyl sulphate-polyacrylamide gel (SDS-PAGE) and blotted onto nitrocellulose membranes. Membranes were blocked at room temperature for 1 hour with TBS 1X-5% BSA. Membranes were then incubated in primary antibody (see Key Resources Table) overnight at 4°C. The membranes were then washed with PBST (PBS1X + 0.05% Tween20; Merck) for three times 5 minutes and incubated with anti-mouse secondary antibody (see Key Resources Table). After washing with PBST for 5 minutes twice and with PBS for 5 minutes once, membrane-bound antibodies were detected by fluorescence with the Odyssey ® Fc Imaging System. Alpha-tubulin (1:5000; Sigma) was used as a loading control for normalization and quantification. Images were analyzed with Image Studio Lite Version 5.2 software.

##### ***Histology and Immunocytochemistry***

Specimens were fixed at the indicated time points with 4% paraformaldehyde (153799, Aname) overnight at 4˚C. Samples were then washed twice with PBS, embedded in paraffin and sectioned into 5 to 10 µm samples. Sections were stained for Hematoxylin and Eosin, Masson`s trichrome and Periodic acid–Schiff. Images were captured using the AF7000 Leica microscope. Quantification of collagen fibres was performed as described elsewhere [67] using ImageJ. Briefly, the color deconvolution plugin was used to generate monochromatic images from Masson's trichrome staining images. The integrated intensity of the green component corresponding to the collagen fibers was then calculated using ImageJ. Immunofluorescence samples were fixed with 4% paraformaldehyde (153799, Aname) for 20 minutes at room temperature, washed trice with PBS and blocked using Tris-buffered saline (TBS) containing 6% donkey serum (S30, Millipore) and 1% Triton X-100 (T8787, Sigma) for 1h at room temperature. Samples were treated overnight at 4 °C with the primary antibodies indicated in Key Resources Table diluted in antibody dilution buffer (TBS solution with 6% donkey serum and 0.5% Triton X-100). Samples were then washed three times with antibody dilution buffer and further incubated during 4h at room temperature with fluorescent conjugated secondary antibodies indicated in Key Resources Table. For detection of LTL positive cells (LTL+), samples were stained with biotinylated

LTL (B-1325, Vector Labs) using a streptavidin/biotin blocking kit (SP-2002, Vector Labs). LTL+ cells were detected using Alexa Fluor 488 conjugated with streptavidin (SA5488, Vector Labs). Nuclei were stained using 4,6-diamidino-2- phenylindole (DAPI; 1:5000, D1306, Life Technologies) for 30min. Samples were mounted in Fluoromount-G (0100-01, Southern Biotech) and visualized using a SP5 Leica microscope or Zeiss LSM780 confocal microscope. For quantification of immunofluorescence images, we used a custom-made MATLAB code. Specifically, the percentage of area occupied by NP+ cells within kidney organoids was calculated by adding the area of NP+ cells divided by the entire area of the organoid.

***Electron Microscopy***

Specimens were fixed with 2.5% glutaraldehyde containing 1% tannic acid in 0.1 M phosphate buffer (PB, pH 7.4). Then samples were post-fixed for 1 hour at 4°C with 1% OsO4 in 0.1 M PB. Epoxy resin embedding was performed after graded ethanol series. Then toluidine blue staining was performed in semithin sections using a light microscope. Finally, ultrathin sections were prepared using an EM UC7 ultramicrotome (Leica Microsystems, Mannheim, Germany) and collected on copper grids. Then 4% uranyl acetate and lead citrate were used for staining. Samples were subsequently analyzed with a JEM 1230 electron microscope (JEOL, UK, Ltd).

***qRT-PCR***

RNA was isolated from cells and kidney organoids using Trizol (Invitrogen). 2 µg RNA was reverse transcribed using the cDNA archival kit (Life Technology), and qRT-PCR was run in the ViiA 7 System (Life Technology) machine using SYBRGreen Master Mix (Applied Biosystem) and gene-specific primers. The data were normalized and analyzed using the ΔΔCt method. The primers sequences used are shown in Key Resources Table.

***Oxygen consumption rate (OCR)***

The measurement of OCR proximal tubular cells was performed using XFe24 extracellular flux analyzer (Seahorse Bioscience) as previously described [68]. Briefly, cells were plated at density of 10,000 cells/well in a Seahorse cell culture microplate. Cellular OCR was measured and normalized to protein quantity in each well. The final concentration for oligomycin, FCCP, rotenone and antimycin at 10µM, 2µM, 5µM and 15µM, respectively.

***scRNA-Seq data analysis***

The computational analysis of the resulting UMI count matrices was performed using the R package Seurat (3.2.1)[69]. Poor quality cells were removed, such as those in the lower and upper 2.5% quantiles for the number of UMIs and detected genes per cell in diabetic samples and those with < 1,000 detected genes per cell in ACE2 samples. For the latter, the upper thresholds were obtained from visual inspection of the UMI and detected genes distribution. In all cases cells with more than 10% of UMIs assigned to mitochondrial genes were discarded and genes expressed in less than 3 cells were removed from the analyses. Each dataset was subjected to normalization, identification of highly variable features and scaling using the *SCTransform* function, regressing out for the number of detected genes, the % of mitochondrial UMIs and cell cycle. All samples were integrated independently to remove batch effects among them and enable downstream comparisons between different conditions. The infection samples at differentiating conditions (11mM) were processed and integrated in the same way as 5mM and high oscillatory glucose conditions.

Principal component analysis was performed, and the top 20 components were kept for further analysis in the integrated ACE2 dataset. In the diabetic data set the top 40 component were kept for analysis. Clustering was performed by setting the resolution parameter to 0.6 in both cases Dimensional reduction for data visualization was done using the RunUMAP function. Cell markers in each cluster were identified using the *FindConservedMarkers* and *FindAllMarkers* functions in the non-integrated counts by using the Wilcoxon Rank Sum test. Genes with p-value < 0.05 (adjusted by Bonferroni's correction) were retained. Clusters were labelled by comparing the expression of the identified markers with publicly available databases located in KIT (Kidney Interactive Transcriptomics) webpage [http://humphreyslab.com/SingleCell/] and with markers from previous publications [70]. Cell type annotations from the diabetic samples were transferred to the 11 mM analysis using the TransferData function in Seurat. Differential expression analysis to identify changes across conditions were also performed using the Wilcoxon test, keeping as up-regulated genes those present in a fraction of 0.1 of either population, with a log fold-change greater than 0.1 and with an adjusted p-value < 0.05. To perform over-representation analysis the R package enrichR [71] was used with the database ¨MSigDB_Hallmark_2020". Gene set enrichment analysis (GSEA) was done with the fgsea R package [72], considering the hallmark gene sets from MSigDB.

***Data Representation and Statistical Analysis***

Student's *t*-test was used to analyze differences between two groups, and One-way or Two-way ANOVA was used to analyze intergroup differences (tukey's multiple comparisons test). *P*-values less than 0.05 were considered statistically significant. The analysis was performed using GraphPad Prism 5 (GraphPad software). Densitometry results of Western Blots were quantified using ImageJ software. All data are presented as mean ± SEM and other details such as the number of replicates and the level of significance is mentioned in figure legends and supplementary tables.

##### *Data and Code Availability*

Raw sequencing data for the single cell kidney organoid reported in this paper were deposited in Gene Gene Expression Omnibus (GEO) under the accession number GEO: GSE181002

##### *Key Resources Table*

| REAGENT or RESOURCE | SOURCE | IDENTIFIER |
|---|---|---|
| **Antibodies** | | |
| **Primary antibodies for Immunoblots** | | |
| Human ACE-2 | Bio-Techne R&D Systems S.L | Cat# AF933-SP |
| Collagen I | Abcam | Cat# ab34710; RRID:AB_731684 |
| α-Tubulin | Abcam | Cat# ab4074; RRID:AB_2288001 |
| **Secondary antibodies for Immunoblots** | | |
| IRDye® 680RD Goat anti-Mouse IgG Secondary Antibody | Licor | Cat# 925-68070; RRID:AB_2651128 |
| IRDye® 800CW Goat anti-Rabbit IgG Secondary Antibody | Licor | Cat# 925-32211; RRID:AB_2651127 |
| **Primary antibodies for Immunofluorescence** | | |
| Human ACE2 | Bio-Techne R&D Systems | Cat# AF933; RRID:AB_355722 |
| Laminin | Merck | Cat# L9393; RRID:AB_477163 |
| Collagen I | Abcam | Cat# ab34710; RRID:AB_731684 |

| | | |
|---|---|---|
| Collagen IV | Merck | Cat# AB769; RRID:AB_92262 |
| Fibronectin | Abcam | Cat# ab2413; RRID:AB_2262874 |
| Lotus Tetragonolobus Lectin (LTL), Biotinylated | Vector laboratories | Cat# B-1325; RRID:AB_2336558 |
| Podocin | Merck | Cat# P0372; RRID:AB_261982 |
| E-Cadherin | BD Bioscience | Cat# 610181; RRID:AB_397580 |
| Human Podocalyxin Biotinylated | R&D Systems | Cat# BAF1658; RRID:AB_356080 |
| SARS-CoV/SARS-CoV-2 Nucleocapsid | Abyntek Biopharma | Cat# 40143-MM05; RRID:AB_2827977 |
| Human KIM-1 | R&D Systems | Cat# AF1750; RRID:AB_2116561 |
| CD147 | Abcam | Cat# ab666; RRID:AB_305632 |
| Human Nephrin | R&D Systems | Cat# AF4269 RRID:AB_2154851 |
| Recombinant PE Anti-Sodium Potassium ATPase | Abcam | Cat# ab209299 |
| PGC1$\alpha$ | R&D Systems | Cat# NBP1-04676 RRID:AB_1522118 |
| **Secondary antibodies for Immunofluorescence** | | |
| Anti-Goat Alexa Fluor 488-conjugated | Jackson ImmunoResea rch | Cat# 705-545-147; RRID:AB_2336933 |
| Anti-Goat IgG Alexa Fluor 555-conjugated | Fisher Scientific | Cat# A-21432; RRID:AB_2535853 |
| Anti-rabbit IgG Alexa fluor 488-conjugated | Fisher Scientific | Cat# A21206; RRID:AB_2535792 |

| | | |
|---|---|---|
| Anti-rabbit IgG Alexa fluor 555-conjugated | Fisher Scientific | Cat# A-31572; RRID:AB_162543 |
| Anti-Mouse IgG CyTM3-conjugated | Jackson ImmunoResearch | Cat# 715-165-151; RRID:AB_2315777 |
| Anti-Sheep IgG Alexa Fluor 555-conjugated | Fisher Scientific | Cat# A-21436; RRID:AB_2535857 |
| Dylight 649 Streptavidin | Vector Labs | Cat# SA-5649; RRID:AB_2336421 |
| **Reagents for FACS** | | |
| Fluorescein-conjugated LTL | Vector Labs | Cat# FL-1321 |
| **Biological Samples** | | |
| Human kidney tubular cells | | N/A |
| **Chemicals, Peptides, and Recombinant Proteins** | | |
| Essential 8 medium | ThermoFisher | Cat# A1517001 |
| Vitronectin | ThermoFisher | Cat# A14700 |
| Trypan blue solution | Sigma | Cat# T8154 |
| EDTA solution | ThermoFisher | Cat# 15575-038 |
| Accumax | Stem Cell Technologies | Cat# 07921 |
| RPMI 1640 | ThermoFisher | Cat#21875-034 |
| Advanced RPMI 1640 | ThermoFisher | Cat#12633020 |
| GlutaMAX (200 mM) | ThermoFisher | Cat# 35050–038 |
| Penicillin/Streptomicin | ThermoFisher | Cat# 15140122 |
| Dulbecco's modified eagle medium (DMEM) glucose-free | ThermoFisher | Cat# 11966025 |
| D-Glucose | Merck | Cat# G7021 |
| D-Manitol | Merck | Cat# M9647 |

| | | |
|---|---|---|
| ITS | Merck | Cat#I3146 |
| Fetal Bovine Serum (FBS) | Gibco | Cat# 10270-106 |
| Human EGF | R&D systems | Cat# 236-EG-01M |
| CHIR99021 | Merck | Cat#SML1046; CAS: 2 52917-06-9 |
| Recombinant human FGF9 | PeproTech | Cat# 100-23 |
| Heparin | Merck | Cat# H3149; CAS: 9041-08-1 |
| Activin A | R&D systems | Cat# 338-AC-050 |
| Cell culture grade distilled water | ThermoFisher | Cat# 15230-089 |
| Protease inhibitor cocktail | Roche | Cat#11836153001 |
| Phosphate buffered saline (PBS) pH 7.4 (1x) | ThermoFisher | Cat#1001–015 |
| RIPA buffer | Cell signaling | Cat#9806 |
| SYBR Green PCR Master Mix | Applied Biosystem | Cat# KK4605 |
| Fluoromount-G | Southern Biotech | Cat# 0100-01 |
| Triton X-100 | Sigma | Cat# T8787 |
| 4',6-Diamidino-2-Phenylindole, Dihydrochloride (DAPI) | ThermoFisher | Cat# D1306 |
| Dimethyl Sulfoxide (DMSO) | Merck | Cat# D2650; CAS: 67-68-5 |
| Paraformaldehyde | Aname | Cat# sc-281692 |
| Donkey serum | Millipore | Cat# S30 |
| Glutaraldehyde | Sigma-Aldrich | Cat# G7776 |
| Sodium phosphate dibasic dihydrate ≥99.0% | Sigma-Aldrich | Cat# 71643 |
| Sodium phosphate monobasic ≥98% | Sigma-Aldrich | Cat# S3139 |
| Trizol | ThermoFisher | Cat# 15596018 |
| Chloroform | Sigma Merck | Cat# 1.02445 CAS:67-66-3 |
| 2-Propanol | Panreac | Cat# 131090 CAS: 67-63-0 |

| | | |
|---|---|---|
| Ethanol | VWR | Cat# 100983.2500<br>CAS: 64-17-5 |
| Actinomycin A | Sigma | Cat# A8674-25MG |
| Nuclease-Free Water | Ambion | Cat# AM9937 |
| Lipofectamine RNAiMAX Transfection Reagent | ThermoFisher | Cat# 13778150 |
| **Critical Commercial Assays** | | |
| BCA Protein Assay Kit | ThermoFisher | Cat# 23225 |
| cDNA Reverse Transcription Kit | Applied Biosystems | Cat# 4368813 |
| Seahorse XFe96 FluxPak mini | Agilent Technologies | Cat# 102601-100 |
| Chromium Single Cell 3’ Library & Gel Bead Kit V3 | 10X Genomics (USA) | Cat# PN-1000075 |
| NSQ 500/550 Hi Output KT v2.5 (75 CYS) | Illumina (San Diego, CA 92122 USA) | Cat# 20024906 |
| Streptavidin/Biotin blocking kit | Vector laboratories | Cat# SP-2002 |
| **Deposited Data** | | |
| scRNA seq data kidney organoids | This study | GEO: GSE181002 |
| **Experimental Models: Organisms/Strains** | | |
| N/A | | |
| **Oligonucleotides** | | |
| **crRNA sequences** | This paper | N/A |
| ACE2ko Cr1 (5’-3’) TAGACTACAATGAGAGGCTC | N/A | N/A |
| ACE2ko Cr2 (5’-3’) GCCATTATATGAAGAGTATG | N/A | N/A |
| BSGko Cr1 (5’-3’) CTTGGAGCCAAGGTCTTCTA | N/A | N/A |

| | | |
|---|---|---|
| BSGko Cr2 (5'-3') TTCACTACCGTAGAAGACCT | N/A | N/A |
| **MiSeq oligos** | This paper | N/A |
| MiSeq_ACE2ko_F1 (5'-3') ACACTCTTTCCCTACACGACGCTCTTCCGATCTTGTG TGCTTTGGGATAACAGGT | N/A | N/A |
| MiSeq_ACE2ko_R1 (5'-3') GACTGGAGTTCAGACGTGTGCTCTTCCGATCTGCCAC ACAGAGAGCTTCAGG | N/A | N/A |
| MiSeq_BSGko_F2 (5'-3') ACACTCTTTCCCTACACGACGCTCTTCCGATCTGGGG AGGAGCCGCAGGTTC | N/A | N/A |
| MiSeq_BSGko_R2 (5'-3') GACTGGAGTTCAGACGTGTGCTCTTCCGATCTCGTCC TCCTTCAGCACCACG | N/A | N/A |
| **qPCR oligos** | This paper | N/A |
| SARS-CoV-2 (oligo1)_Forward GCCTCTTCTCGTTCCTCATCAC | Eurofins | N/A |
| SARS-CoV-2 (oligo1)_Reverse AGCAGCATCACCGCCATTG | Eurofins | N/A |
| SARS-CoV-2 (oligo2)_Forward AGCCTCTTCTCGTTCCTCATCAC | Eurofins | N/A |
| SARS-CoV-2 (oligo2)_Reverse CCGCCATTGCCAGCCATTC | Eurofins | N/A |
| TMPRSS2_Forward GTCCCCACTGTCTACGAGGT | Thermo Fisher | N/A |
| TMPRSS2_Reverse CAGACGACGGGGTTGGAAG | Thermo Fisher | N/A |
| NRP1_Forward GGCGCTTTTCGCAACGATAAA | Thermo Fisher | N/A |
| NRP1_Reverse TCGCATTTTTCACTTGGGTGAT | Thermo Fisher | N/A |
| BSG_Forward CCGCAACCACCTTACTCG | Thermo Fisher | N/A |
| BSG_Reverse GGACAGAGGTTTGGATGGTG | Thermo Fisher | N/A |
| ACE2_Forward CGAAGCCGAAGACCTGTTCTA | Thermo Fisher | N/A |

| | | |
|---|---|---|
| ACE2_Reverse GGGCAAGTGTGGACTGTTCC | Thermo Fisher | N/A |
| HK2_Forward AGCCCTTTCTCCATCTCCTT | Thermo Fisher | N/A |
| HK2_Reverse AACCATGACCAAGTGCAGAA | Thermo Fisher | N/A |
| LDHA_Forward GGAGATCCATCATCTCTCCC | Thermo Fisher | N/A |
| LDHA_Reverse GGCCTGTGCCATCAGTATCT | Thermo Fisher | N/A |
| PGC1α _Forward CTGCTAGCAAGTTTGCCTCA | Thermo Fisher | N/A |
| PGC1α _Reverse AGTGGTGCAGTGACCAATCA | Thermo Fisher | N/A |
| CD36_Forward TCAATTCGTCTAATCATTGGAAA | Thermo Fisher | N/A |
| CD36_Reverse GCAAGACTCTGGAGCCAGTC | Thermo Fisher | N/A |
| COL3A1_Forward AGGACTGACCAAGATGGGAA | Thermo Fisher | N/A |
| COL3A1_Reverse AGGGGAGCTGGCTACTTCTC | Thermo Fisher | N/A |
| COL4A1_Forward CCTTTTGTCCCTTCACTCCA | Thermo Fisher | N/A |
| COL4A1_Reverse CTCCACGAGGAGCACAGC | Thermo Fisher | N/A |
| RPLP0 _Forward CCATTCTATCATCAACGGGTACAA | Thermo Fisher | N/A |
| RPLP0 _Reverse AGCAAGTGGGAAGGTGTAATCC | Thermo Fisher | N/A |
| SLC16A1_Forward GGCTGTCATGTATGGTGGAG | Thermo Fisher | N/A |
| SLC16A1_Reverse GACAAGCAGCCACCAACAATC | Thermo Fisher | N/A |
| PODXL_Forward GATAAGTGCGGCATACGGCT | Thermo Fisher | N/A |
| PODXL_Reverse GCTCGTACACATCCTTGGCA | Thermo Fisher | N/A |
| WT1_Forward GCCAGGATGTTTCCTAACGC | Thermo Fisher | N/A |
| WT1_Reverse CGAAGGTGACCGTGCTGTAA | Thermo Fisher | N/A |
| NPHS1_Forward GGCTCCCAGCAGAAACTCTT | Thermo Fisher | N/A |
| NPHS1_Reverse CACAGACCAGCAACTGCCTA | Thermo Fisher | N/A |
| SLC3A1_Forward CACCAATGCAGTGGGACAAT | Thermo Fisher | N/A |
| SLC3A1_Reverse CTGGGCTGAGTCTTTTGGAC | Thermo Fisher | N/A |

| | | |
|---|---|---|
| SLC12A1_Forward AGTGCCCAGTAATACCAATCGC | Thermo Fisher | N/A |
| SLC12A1_Reverse GCCTAAAGCTGATTCTGAGTCTT | Thermo Fisher | N/A |
| SLC12A3_Forward CCTGGGTGGAGACCTTCATTC | Thermo Fisher | N/A |
| SLC12A3_Reverse GAGCCCCAATTTACCTCTGGC | Thermo Fisher | N/A |
| CDH16_Forward AGAATGACAACGTGCCTATCTG | Thermo Fisher | N/A |
| CDH16_Reverse GCTGACAGTCTAGTCACTTCAGT | Thermo Fisher | N/A |
| MAFB_Forward GACGCAGCTCATTCAGCAG | Thermo Fisher | N/A |
| MAFB_Reverse CTCGCACTTGACCTTGTAGGC | Thermo Fisher | N/A |
| **Recombinant DNA** | | |
| N/A | | |
| **Experimental Models: Cell Lines** | | |
| ES[4] Human Embryonic Stem Cell line | The National Bank of Stem Cells (ISCIII,Madrid) | https://www.isciii.es/ |
| CBiPS1sv-4F-40 | The National Bank of Stem Cells (ISCIII,Madrid) | https://www.isciii.es/ |
| Vero E6 | ATCC CRL 1586 | |
| **Software and Algorithms** | | |
| ImageJ | NIH | https://imagej.nih.gov/ij |
| Prism 5 | Graphpad Software | https://www.graphpad.com/scientific-software/prism |
| Image Studio Lite Version 5.2 software | LICOR | https://www.licor.com/bio/image-studio-lite/d5 |
| FlowJo Software | FlowJo | N/A |

| | | |
|---|---|---|
| Cell Ranger 4.0.0 | 10x Genomics | https://support.10xgenomics.com/single-cell-gene-expression/software/downloads/latest |
| Seurat R package 3.2.1 | open source | https://satijalab.org/seurat/ |
| fgsea R package 1.16.0 | open source | http://bioconductor.org/packages/release/bioc/html/fgsea.html |
| enrichR R package v3.0 | open source | https://CRAN.R-project.org/package=enrichR |

#### Supplemental Figures and Legends

### Figure S1

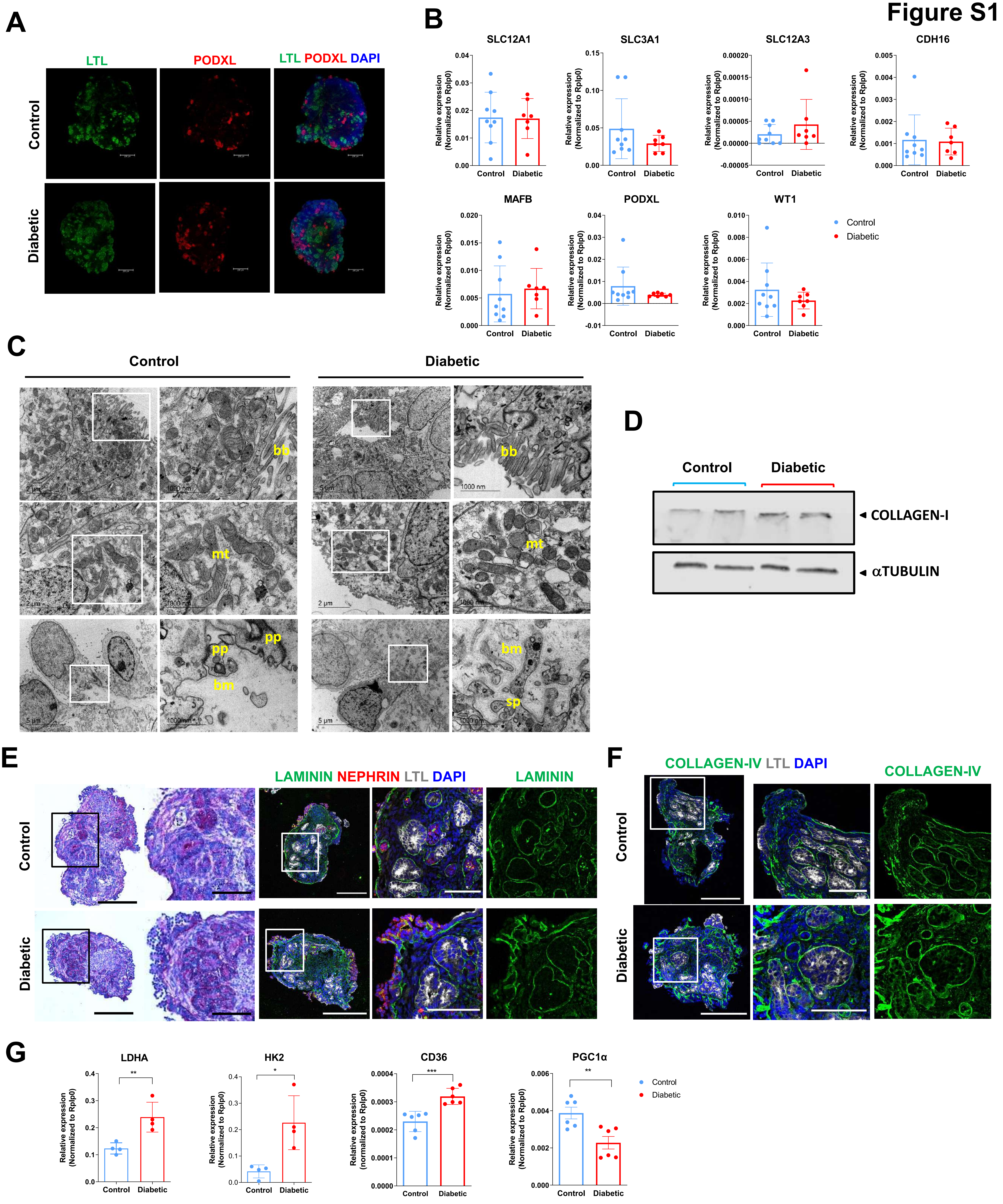

A
LTL
PODXL
LTL PODXL DAPI
Control
Diabetic
B
SLC12A1
SLC3A1
SLC12A3
CDH16
MAFB
PODXL
WT1
Relative expression (Normalized to Rplp0)
Control
Diabetic
C
Control
Diabetic
bb
mt
pp
bm
sp
2 µm
1000 nm
5 µm
D
Control
Diabetic
COLLAGEN-I
αTUBULIN
E
LAMININ NEPHRIN LTL DAPI
LAMININ
Control
Diabetic
F
COLLAGEN-IV LTL DAPI
COLLAGEN-IV
Control
Diabetic
G
LDHA
HK2
CD36
PGC1α
Relative expression (Normalized to Rplp0)
Control
Diabetic
**
*
***
**

**Figure S1. High Oscillatory Glucose Regime Promotes Diabetogenic-like Responses in Human Kidney Organoids, Related to Figure 1.**

A. Representative immunofluorescence staining of LTL (green), Podocalyxin (PODXL; red) and DAPI (blue) in kidney organoids cultured under Control or Diabetic conditions for 7 days. Scale bars, 250 μm.

B. mRNA expression level of proximal tubule markers (SLC12A1, SLC3A1, SLC12A4, CDH16) and podocyte markers (MAFB, PODXL, WT1) in Control and Diabetic kidney organoids. The data are represented as mean ± SEM. N>7 independent experimental replicates from a pool of 12 organoids/group; paired t-test.

C. TEM in kidney organoids cultured in Control or Diabetic conditions for 7 days. Magnified views of the boxed regions show details of tubular-like epithelial cell-related structures including brush borders (bb) and high mitochondrial (mit) content (upper and middle panels). Bottom panels show podocyte-related structures, including the deposition of a basement membrane (bm), and primary (pp) and secondary cell processes (sp). Scale bars, 2 μm and 5 μm; 1000 nm (magnified views).

D. Protein levels of COLLAGEN-I in kidney organoids exposed to Control Diabetic conditions are shown by Western Blot. aTUBULIN was used as loading control. *n* = 2 independent experimental replicates from a pool of 12 organoids/group.

E. Representative Periodic Acid-Schiff (PAS) staining in Control and Diabetic kidney organoids. Scale bars, 250 μm, 100 μm (magnified views). Consecutive sections were stained for LAMININ (green), NEPHRIN (red), LTL (grey) and DAPI (blue). Scale bars, 250 μm, 100 μm (magnified views).

F. Representative immunofluorescence staining of COLLAGEN-IV (green) LTL (grey) and DAPI (blue) in Control and Diabetic kidney organoids. Scale bars, 250 μm, 100 μm (magnified views).

G. mRNA expression level of glycolytic enzymes (LDHA and HK2) and FAO markers (CD36 and PGC1a) in Control and Diabetic kidney organoids. Data are represented as mean ± SEM. N>4 independent experimental replicates from a pool of 12 organoids/group; *P < 0.05, **P < 0.01 ***P < 0.001, paired Student's t-test.

Figure S2

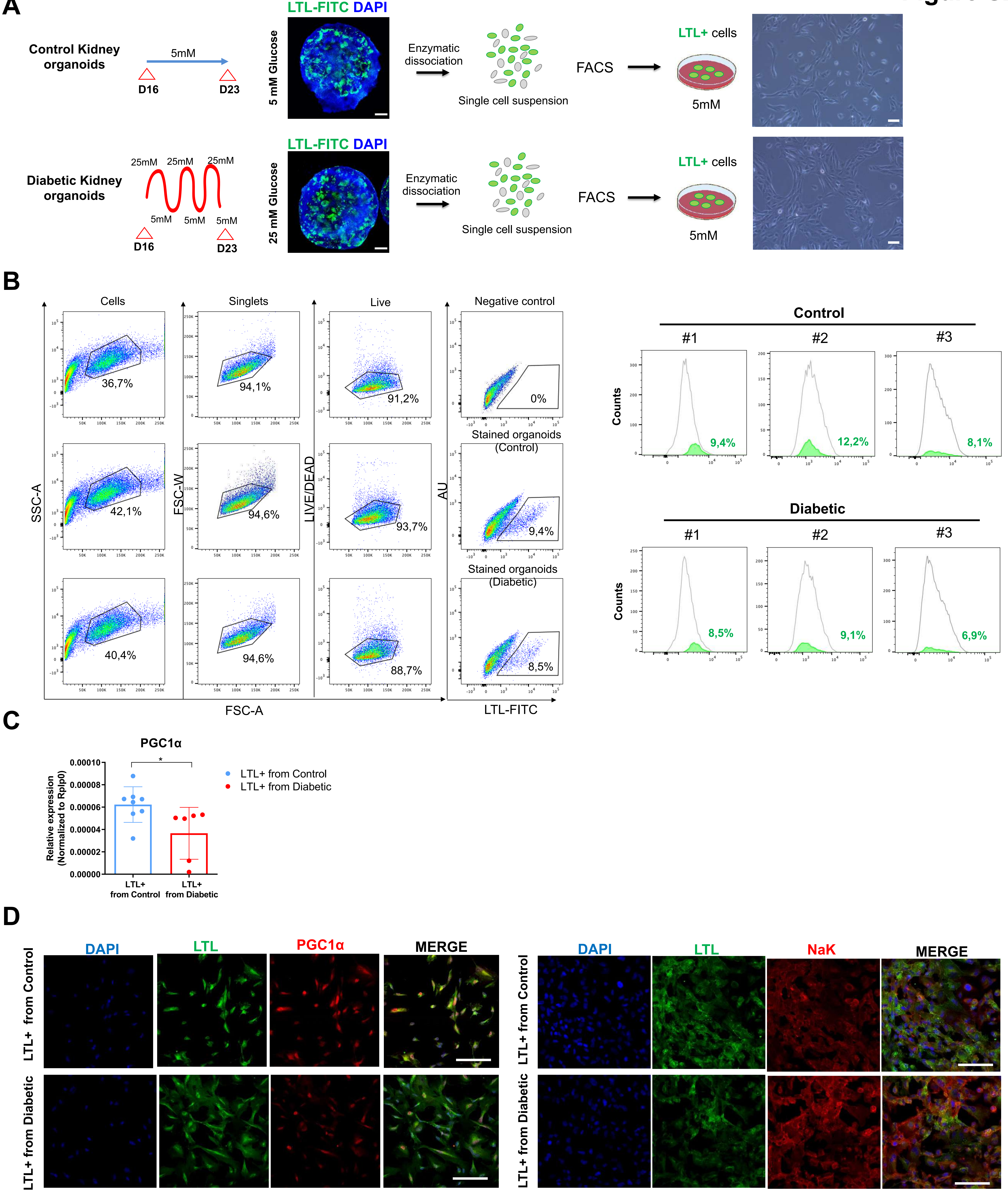
A
Control Kidney organoids
5mM
D16
D23
LTL-FITC DAPI
5 mM Glucose
Enzymatic dissociation
Single cell suspension
FACS
LTL+ cells
5mM
Diabetic Kidney organoids
25mM 25mM 25mM
5mM 5mM 5mM
D16
D23
LTL-FITC DAPI
25 mM Glucose
Enzymatic dissociation
Single cell suspension
FACS
LTL+ cells
5mM
B
Cells
Singlets
Live
Negative control
36,7%
94,1%
91,2%
0%
Stained organoids (Control)
42,1%
94,6%
93,7%
9,4%
Stained organoids (Diabetic)
40,4%
94,6%
88,7%
8,5%
SSC-A
FSC-W
LIVE/DEAD
AU
FSC-A
LTL-FITC
Control
#1 #2 #3
Counts
9,4%
12,2%
8,1%
Diabetic
#1 #2 #3
Counts
8,5%
9,1%
6,9%
C
PGC1α
Relative expression (Normalized to Rplp0)
0.00010
0.00008
0.00006
0.00004
0.00002
0.00000
*
LTL+ from Control
LTL+ from Diabetic
LTL+ from Control
LTL+ from Diabetic
D
DAPI
LTL
PGC1α
MERGE
LTL+ from Control
LTL+ from Diabetic
DAPI
LTL
NaK
MERGE
LTL+ from Control
LTL+ from Diabetic

**Figure S2. Characterization of Tubular Epithelial Cells Derived from Diabetic Human Kidney Organoids Reveals Alterations in Cellular Metabolism, Related to Figure 1.**

A. Kidney organoids exposed to Control or Diabetic conditions were stained with LTL-FITC and dissociated to single cells. Proximal-like epithelial cells (LTL+ cells) were isolated by cell sorting for LTL marker expression and further cultured in 5 mM glucose culture medium. Right panels show representative bright field images. Scale bars, 100 μm.

B. Gating strategy for cell sorting of LTL+ cells from kidney organoids exposed to Control or Diabetic conditions. Numbers in outlined areas indicate percent cells. AU, autofluorescence. Histograms representing the overlay of live cells and LTL+ cells isolated from Control and Diabetic kidney organoids from 3 independent cell sorting experiments.

C. mRNA expression levels of PGC1a in LTL+ cells isolated from kidney organoids exposed to Control or Diabetic conditions. Mean ±SEM is calculated from N>6 biological replicates/ experimental condition.

D. Representative immunofluorescence staining in LTL+ cells isolated from Control or Diabetic kidney organoids and subsequently expanded in 5 mM glucose culture medium for the expression of proximal tubular cell markers including LTL (green), PGC1a (red) and Na-K ATPase (NaK; red). Nuclei were counterstained with DAPI (blue). Scale bars, 100 μm.

### Figure S3

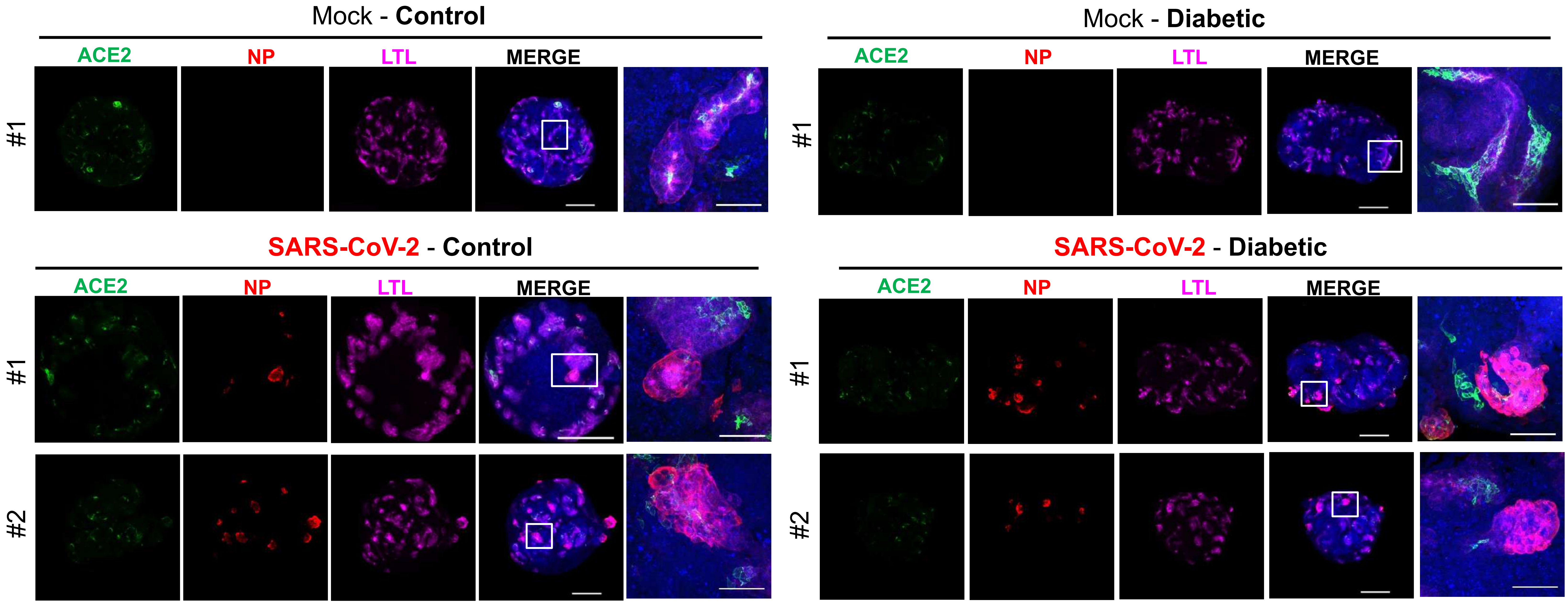
Mock - Control
ACE2
NP
LTL
MERGE
#1
Mock - Diabetic
ACE2
NP
LTL
MERGE
#1
SARS-CoV-2 - Control
ACE2
NP
LTL
MERGE
#1
#2
SARS-CoV-2 - Diabetic
ACE2
NP
LTL
MERGE
#1
#2

**Figure S3. SARS-CoV-2 Infection in Diabetic Human Kidney Organoids, Related to Figure 3.**
Immunofluorescence staining of Mock or SARS-CoV-2 infected ($10^6$ virus particles/organoid as determined in Vero cells) Control or Diabetic kidney organoids at 1 day post infection (1 dpi) for the detection of ACE2 (green), virus nuclear protein (NP; red), LTL (magenta) and DAPI (blue). Scale bars, 250 μm, 50 μm (magnified views). *n = 1* organoid (Mock Control; Mock Diabetic). *n = 2* organoids (SARS-CoV-2 Control; SARS-CoV-2 Diabetic.

### Figure S4

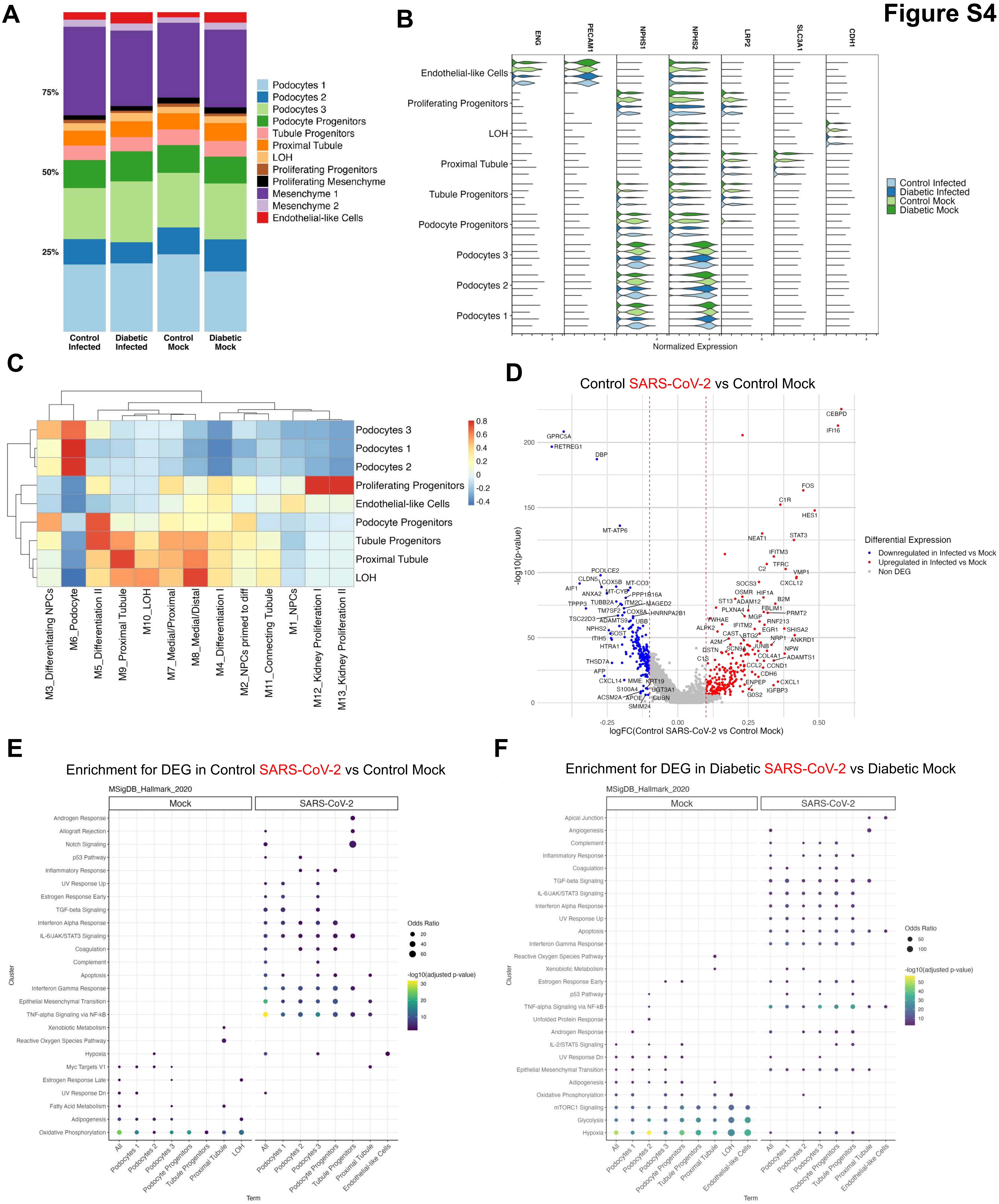
A
B
C
D
E
F
Control Infected
Diabetic Infected
Control Mock
Diabetic Mock
Normalized Expression
Control SARS-CoV-2 vs Control Mock
Differential Expression
Downregulated in Infected vs Mock
Upregulated in Infected vs Mock
Non DEG
logFC(Control SARS-CoV-2 vs Control Mock)
-log10(p-value)
Enrichment for DEG in Control SARS-CoV-2 vs Control Mock
Enrichment for DEG in Diabetic SARS-CoV-2 vs Diabetic Mock
MSigDB_Hallmark_2020
Mock
SARS-CoV-2
Odds Ratio
-log10(adjusted p-value)
Cluster
Term

**Figure S4. Phenotypic and Transcriptional Changes in Diabetic Human Kidney Organoids after SARS-CoV-2 Infection, Related to Figure 3, Figure 4.**

A. Cell type proportions in Mock or SARS-CoV-2 infected ($10^6$ virus particles/organoid as determined in Vero cells) kidney organoids exposed to Control or Diabetic conditions after discarding non-renal cells and Proliferating Progenitors 2 clusters.
B. Violin plots for the indicated markers in Mock or SARS-CoV-2 infected ($10^6$ virus particles/organoid as determined in Vero cells) kidney organoids exposed to Control or Diabetic conditions.
C. Correlation analysis between scores for gene modules from second semester fetal-kidney cell types [73] and scores for gene signatures from the annotated renal-like cell types in Mock and SARS-CoV-2 infected kidney organoids. Scores were obtained with the *AddModuleScore* function from Seurat. Color scale denotes Pearson correlation coefficients.
D. Differentially expressed genes (DEGs) in the comparison of SARS-CoV-2 infected ($10^6$ virus particles/organoid as determined in Vero cells) against Mock kidney organoids in Control conditions considering only renal-like cell types (Podocytes 1/2/3, PT, LOH, Tubule Progenitors, Podocyte Progenitors, Proliferating Progenitors and Endothelial-like Cells). In the volcano plot, the x-axis indicates log fold change (FC) and the y-axis indicates statistical significance with the –log10(p-value). Genes with an adjusted p-value < 0.05 are considered upregulated (red) if the logFC > 0.1 and downregulated (blue) if the logFC < -0.1. Non DEG are shown in grey.
E. Gene over-representation analysis using the Hallmark database considering DEGs (adjusted p-value < 0.05 and logFC > 0.1) to compare Mock and SARS-CoV-2 infected ($10^6$ virus particles/organoid as determined in Vero cells) kidney organoids cultured in Control conditions. Each column corresponds to the analysis of DEG upregulated in each condition. Only the ten gene sets with lowest adjusted p-value in each comparison are shown. Circles are coded by color (p-value) and size (Odds ratio).
F. Gene over-representation analysis using the Hallmark database considering DEGs (adjusted p-value < 0.05 and logFC > 0.1) to compare Mock and SARS-CoV-2 infected ($10^6$ virus particles/organoid as determined in Vero cells) kidney organoids cultured in Diabetic conditions. Each column corresponds to the analysis of DEG upregulated in each condition. Only the ten gene sets with lowest adjusted p-value in each comparison are shown. Circles are coded by color (p-value) and size (Odds ratio).

#### Figure S5

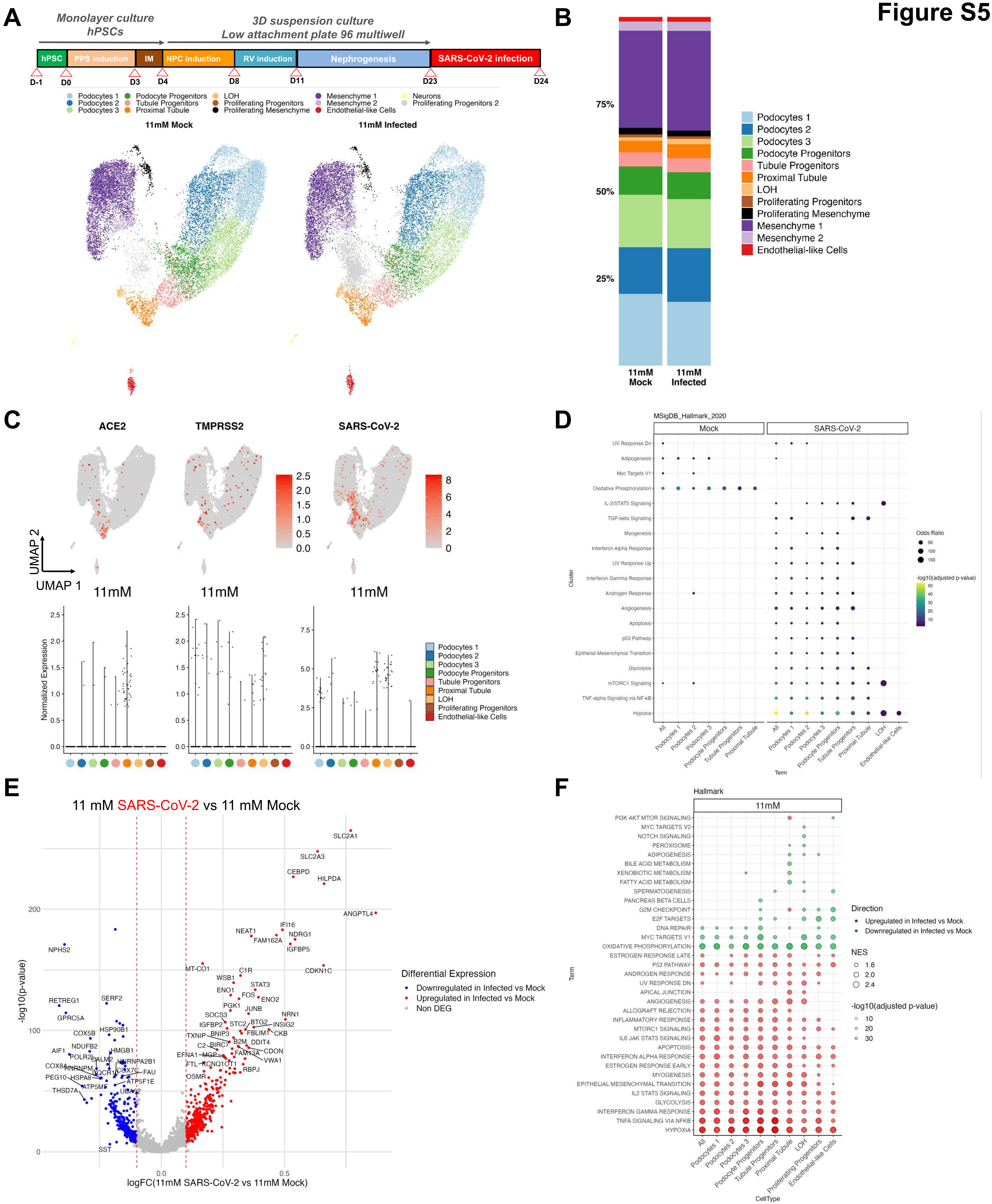

A
Monolayer culture hPSCs
3D suspension culture
Low attachment plate 96 multiwell
hPSC
PPS induction
IM
NPC induction
RV induction
Nephrogenesis
SARS-CoV-2 infection
D-1
D0
D3
D4
D8
D11
D23
D24
11mM Mock
11mM Infected
B
11mM Mock
11mM Infected
Podocytes 1
Podocytes 2
Podocytes 3
Podocyte Progenitors
Tubule Progenitors
Proximal Tubule
LOH
Proliferating Progenitors
Proliferating Mesenchyme
Mesenchyme 1
Mesenchyme 2
Endothelial-like Cells
C
ACE2
TMPRSS2
SARS-CoV-2
UMAP 1
UMAP 2
Normalized Expression
D
MSigDB_Hallmark_2020
Mock
SARS-CoV-2
Odds Ratio
-log10(adjusted p-value)
E
11 mM SARS-CoV-2 vs 11 mM Mock
Differential Expression
Downregulated in Infected vs Mock
Upregulated in Infected vs Mock
Non DEG
logFC(11mM SARS-CoV-2 vs 11mM Mock)
-log10(p-value)
F
Hallmark
11mM
Direction
Upregulated in Infected vs Mock
Downregulated in Infected vs Mock
NES
-log10(adjusted p-value)
CellType
Term

**Figure S5. Single Cell RNA Sequencing Analysis of SARS-CoV-2 Infected Human Kidney Organoids exposed to 11 mM Glucose Conditions, Related to Figure 3, Figure 4.**

A. Schematics for the infection of kidney organoids under 11 mM glucose conditions. Uniform manifold approximation and projection (UMAP) of kidney organoids at 1 dpi with SARS-CoV-2 ($10^6$ virus particles/organoid as determined in Vero cells). Clusters are colored by cell type annotation.
B. Cell type proportions in the indicated kidney organoid samples after discarding non-renal cells and Proliferating Progenitors 2 clusters.
C. UMAPs for ACE2, TMPRSS2 and SARS-CoV-2 expression in kidney organoids exposed to 11 mM glucose at 1dpi. Cells are colored based on expression level. For SARS-CoV-2, expression is considered as zero for cells expressing < 5 UMIs.
D. Gene over-representation analysis using the Hallmark database considering DEGs (adjusted p-value < 0.05 and logFC > 0.1) to compare Mock and SARS-CoV-2 infected kidney organoids ($10^6$ virus particles/organoid as determined in Vero cells) cultured in 11 mM glucose. Each column corresponds to the analysis of DEG upregulated in each condition. Only the ten gene sets with lowest adjusted p-value in each comparison are shown. Circles are coded by color (p-value) and size (Odds ratio).
E. Differentially expressed genes (DEGs) in SARS-CoV-2 infected ($10^6$ virus particles/organoid as determined in Vero cells) against Mock samples in 11 mM glucose considering only renal-like cell types (Podocytes 1/2/3, PT, LOH, Tubule Progenitors, Podocyte Progenitors, Proliferating Progenitors y Endothelial-like cells). In the volcano plot, the x-axis indicates log fold change (FC) and the y-axis indicates statistical significance with the –log10(p-value Genes with an adjusted p-value < 0.05 are considered upregulated (red) if the logFC > 0.1 and downregulated (blue) if the logFC < -0.1. Non DEG are shown in grey.
F. A hallmark gene set enrichment analysis (GSEA) was performed for kidney organoids cultured in 11 mM glucose comparing SARS-CoV-2 infected ($10^6$ virus particles/organoid as determined in Vero cells) versus Mock samples. The ten gene sets per direction and sample with lowest adjusted p-value are shown. Each column corresponds to one of the comparisons. Circles are coded by color (direction), size (NES) and transparency (p-value).

### Figure S6

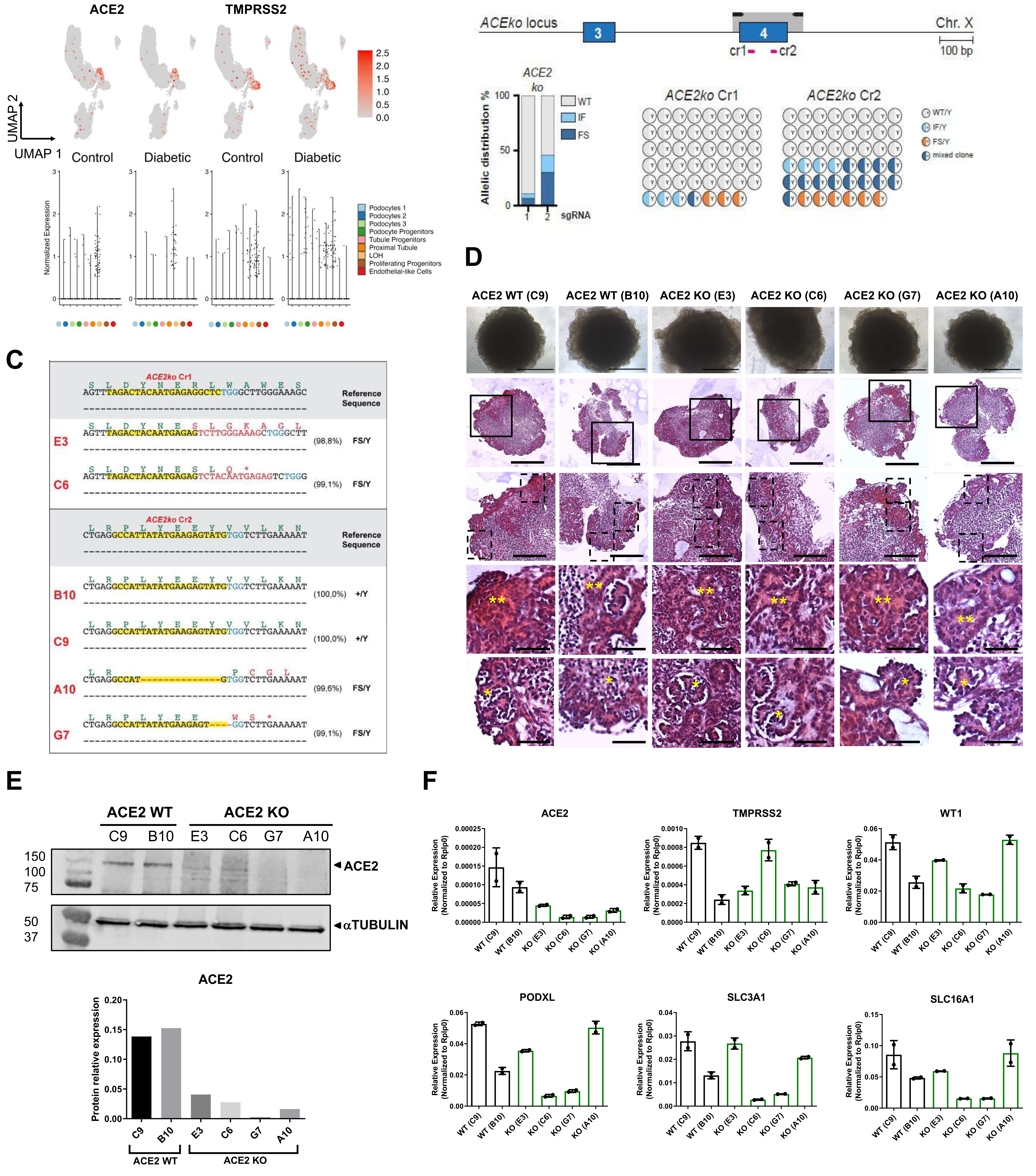

A
ACE2
TMPRSS2
UMAP 2
UMAP 1
Control
Diabetic
Control
Diabetic
2.5
2.0
1.5
1.0
0.5
0.0
Normalized Expression
Podocytes 1
Podocytes 2
Podocytes 3
Podocyte Progenitors
Tubule Progenitors
Proximal Tubule
LOH
Proliferating Progenitors
Endothelial-like Cells
B
ACEko locus
3
4
Chr. X
cr1
cr2
100 bp
ACE2 ko
Allelic distribution %
WT
IF
FS
sgRNA
ACE2ko Cr1
ACE2ko Cr2
WT/Y
IF/Y
FS/Y
mixed clone
C
ACE2ko Cr1
Reference Sequence
E3
(98,8%)
FS/Y
C6
(99,1%)
FS/Y
ACE2ko Cr2
Reference Sequence
B10
(100,0%)
+/Y
C9
(100,0%)
+/Y
A10
(99,6%)
FS/Y
G7
(99,1%)
FS/Y
D
ACE2 WT (C9)
ACE2 WT (B10)
ACE2 KO (E3)
ACE2 KO (C6)
ACE2 KO (G7)
ACE2 KO (A10)
E
ACE2 WT
ACE2 KO
C9
B10
E3
C6
G7
A10
150
100
75
50
37
◄ACE2
◄αTUBULIN
ACE2
Protein relative expression
F
ACE2
TMPRSS2
WT1
PODXL
SLC3A1
SLC16A1
Relative Expression (Normalized to Rplp0)
WT (C9)
WT (B10)
KO (E3)
KO (C6)
KO (G7)
KO (A10)

**Figure S6. Generation and Infection of ACE2 KO Human Kidney Organoids, Related to Figure 5.**

A. UMAPs for ACE2 and TMPRSS2 expression in kidney organoids exposed to Control or Diabetic conditions at 1 dpi. Cells are colored based on expression level. The violin plots in the bottom panels represent expression level for the different cell types indicated in the legend. For SARS-CoV-2, expression is considered as zero for cells expressing < 5 UMIs.
B. Schematic of Cas9/gRNA-targeting sites (pink arrows) in ACE2 *locus* showing exon structure (blue boxes) and PCR amplicons (light gray boxes in all figures in this study). Histogram shows allelic sequence distribution after the transfection of the different gRNAs in undifferentiated human pluripotent stem cells expressing an inducible Cas9 (iCas9). wt, wild-type; mut, mutation; FS, frameshift.
C. Representative sequence of the wild type (+Y) or ACE2 mutant clones generated with the different gRNAs.
D. Representative phase contrast images of ACE2 WT and ACE2 KO kidney organoids. Scale bars, 250 μm. Same specimens were analyzed by Hematoxylin and Eosin staining showing tubular-like (**) and glomerular-like (*) structures. Scale bars, Scale bars, 250 μm, 100 μm (magnified views); 50 μm (magnified views from dashed line boxes).
E. Protein levels of ACE2 in the generated clones are shown by Western Blot. aTUBULIN was used as loading control. Quantification of changes in ACE2 protein expression is shown. Data are represented from a pool of 16 organoids/group.
F. mRNA expression level of ACE2 and TMPRSS2, podocyte markers (WT1 and PODXL) and proximal tubular markers (SLC3A1 and SLC16A1) in ACE2 WT and ACE2 KO kidney organoids. Data from a pool of 12 organoids/group is shown.

Figure S7

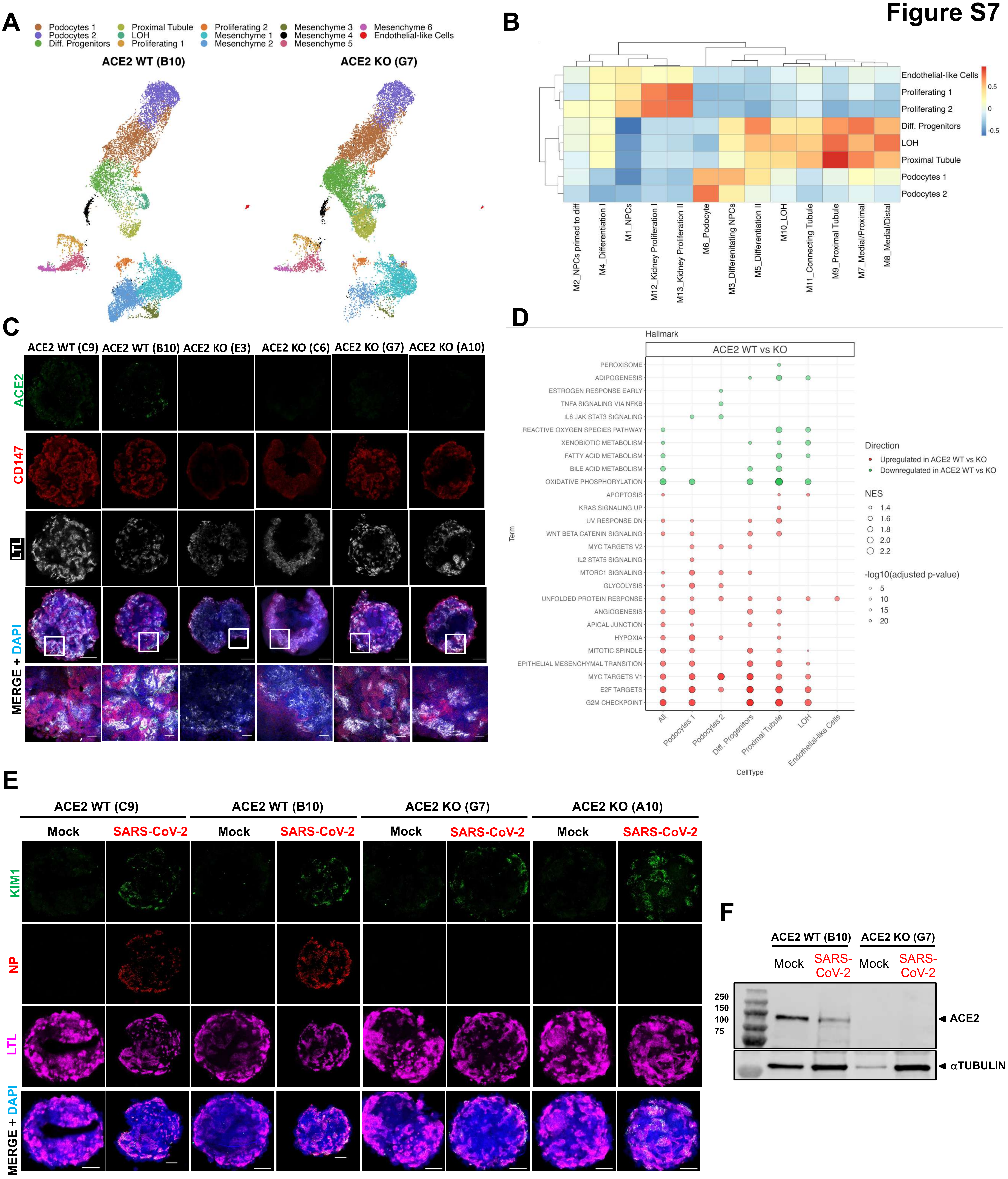

A
Podocytes 1
Podocytes 2
Diff. Progenitors
Proximal Tubule
LOH
Proliferating 1
Proliferating 2
Mesenchyme 1
Mesenchyme 2
Mesenchyme 3
Mesenchyme 4
Mesenchyme 5
Mesenchyme 6
Endothelial-like Cells
ACE2 WT (B10)
ACE2 KO (G7)
B
Endothelial-like Cells
Proliferating 1
Proliferating 2
Diff. Progenitors
LOH
Proximal Tubule
Podocytes 1
Podocytes 2
0.5
0
-0.5
M2_NPCs primed to diff
M4_Differentiation I
M1_NPCs
M12_Kidney Proliferation I
M13_Kidney Proliferation II
M6_Podocyte
M3_Differenitating NPCs
M5_Differentiation II
M10_LOH
M11_Connecting Tubule
M9_Proximal Tubule
M7_Medial/Proximal
M8_Medial/Distal
C
ACE2 WT (C9)
ACE2 WT (B10)
ACE2 KO (E3)
ACE2 KO (C6)
ACE2 KO (G7)
ACE2 KO (A10)
ACE2
CD147
LTL
MERGE + DAPI
D
Hallmark
ACE2 WT vs KO
PEROXISOME
ADIPOGENESIS
ESTROGEN RESPONSE EARLY
TNFA SIGNALING VIA NFKB
IL6 JAK STAT3 SIGNALING
REACTIVE OXYGEN SPECIES PATHWAY
XENOBIOTIC METABOLISM
FATTY ACID METABOLISM
BILE ACID METABOLISM
OXIDATIVE PHOSPHORYLATION
APOPTOSIS
KRAS SIGNALING UP
UV RESPONSE DN
WNT BETA CATENIN SIGNALING
MYC TARGETS V2
IL2 STAT5 SIGNALING
MTORC1 SIGNALING
GLYCOLYSIS
UNFOLDED PROTEIN RESPONSE
ANGIOGENESIS
APICAL JUNCTION
HYPOXIA
MITOTIC SPINDLE
EPITHELIAL MESENCHYMAL TRANSITION
MYC TARGETS V1
E2F TARGETS
G2M CHECKPOINT
Term
All
Podocytes 1
Podocytes 2
Diff. Progenitors
Proximal Tubule
LOH
Endothelial-like Cells
CellType
Direction
Upregulated in ACE2 WT vs KO
Downregulated in ACE2 WT vs KO
NES
1.4
1.6
1.8
2.0
2.2
-log10(adjusted p-value)
5
10
15
20
E
ACE2 WT (C9)
ACE2 WT (B10)
ACE2 KO (G7)
ACE2 KO (A10)
Mock
SARS-CoV-2
KIM1
NP
LTL
MERGE + DAPI
F
ACE2 WT (B10)
ACE2 KO (G7)
Mock
SARS-CoV-2
250
150
100
75
ACE2
αTUBULIN

**Figure S7. SARS-CoV-2 Infection in ACE2 KO Human Kidney Organoids, Related to Figure 5.**

A. UMAP of ACE2 WT kidney organoids (generated from clone B10) and ACE2 KO kidney organoids (generated from ACE2 KO clone G7) colored by annotated cell types.
B. Correlation analysis between scores for gene modules from second semester fetal-kidney cell types [73] and scores for gene signatures from the annotated renal-like cell types in ACE2 WT and ACE2 KO kidney organoids. Scores were obtained with the *AddModuleScore* function from Seurat. Color scale denotes Pearson correlation coefficients.
C. Immunofluorescence staining for ACE2 (green), CD147 (red), LTL (grey) and DAPI (blue) in ACE2 WT and ACE2 KO kidney organoids. Scale bars, 250 µm, 50 µm (magnified views).
D. Hallmark gene set enrichment analysis (GSEA) was performed in kidney organoids generated from ACE2 WT and ACE2 KO lines. The ten gene sets per direction and sample with lowest adjusted p-value are shown. Circles are coded by color (direction), size (NES) and transparency (p-value).
E. Immunofluorescence was performed at 3 dpi in Mock or SARS-CoV-2 infected ($10^6$ virus particles/organoid as determined in Vero cells) ACE2 WT and ACE2 KO kidney organoids. Representative images show the detection of KIM-1 (green), virus nuclear protein (NP; red), LTL (magenta) and DAPI (blue). Scale bars, 250 µm.
F. Protein levels of ACE2 in Mock and SARS-CoV-2 infected ($10^6$ virus particles/organoid as determined in Vero cells) ACE2 WT and ACE2 KO kidney organoids are shown by Western Blot analysis. aTUBULIN was used as loading control. Data from a pool of 12 organoids/group is shown.

**Figure S8**

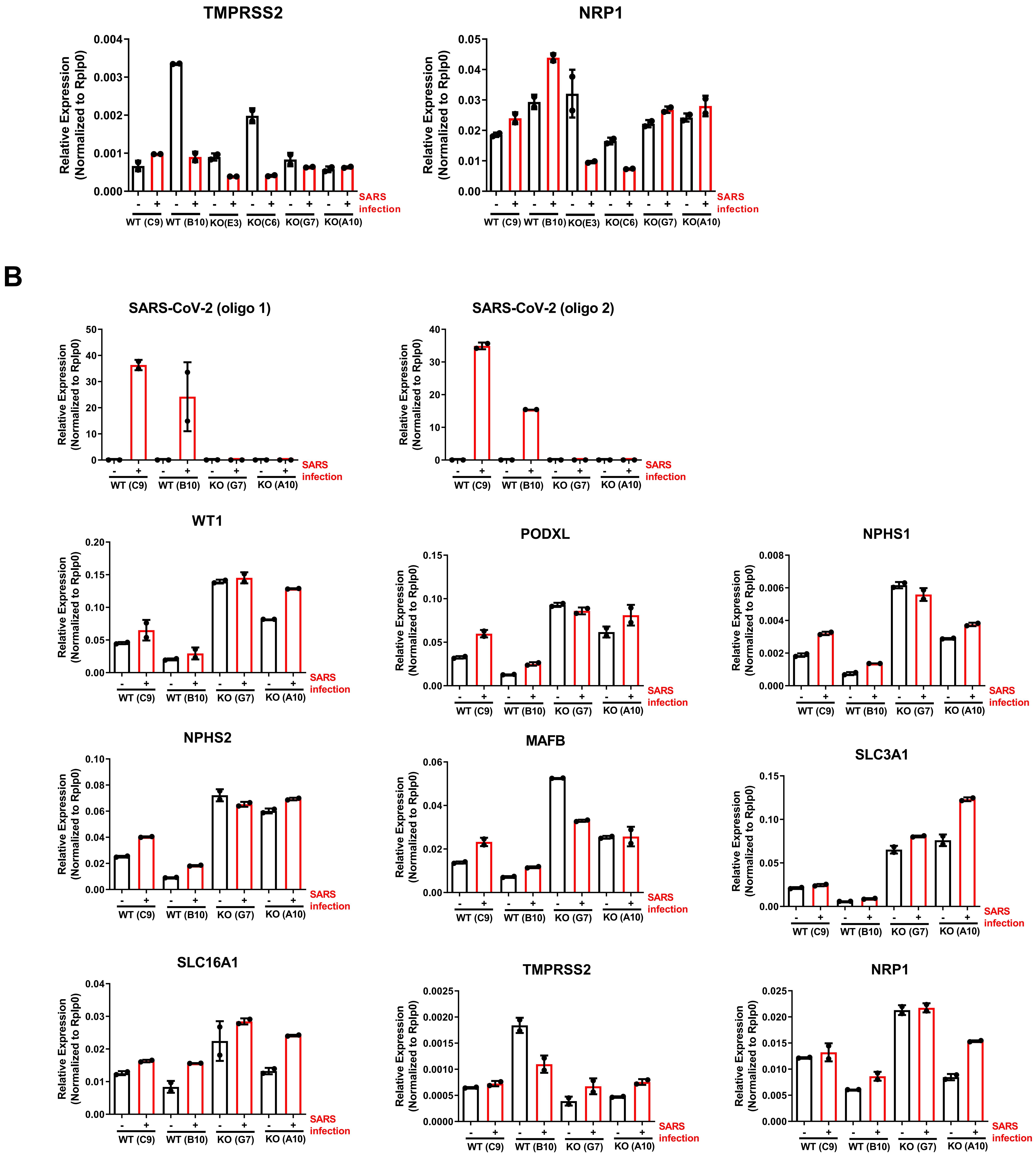
A
TMPRSS2
NRP1
Relative Expression
(Normalized to Rplp0)
SARS
infection
WT (C9)
WT (B10)
KO(E3)
KO(C6)
KO(G7)
KO(A10)
B
SARS-CoV-2 (oligo 1)
SARS-CoV-2 (oligo 2)
WT1
PODXL
NPHS1
NPHS2
MAFB
SLC3A1
SLC16A1
TMPRSS2
NRP1
WT (C9)
WT (B10)
KO (G7)
KO (A10)

**Figure S8. Transcriptional changes upon SARS-CoV-2 Infection in ACE2 WT and KO Human Kidney Organoids, Related to Figure 5.**

A. Quantitative real time was performed at 3 dpi in Mock or SARS-CoV-2 infected ($10^6$ virus particles/organoid as determined in Vero cells) ACE2 WT and ACE2 KO kidney organoids to quantify the mRNA expression levels of TMPRSS2 and NRP1. Data from a pool of 12 organoids/group is shown.

B. Quantitative real time was performed at 3 dpi in Mock or SARS-CoV-2 infected ($10^6$ virus particles/organoid as determined in Vero cells) ACE2 WT and ACE2 KO kidney organoids to quantify the mRNA expression levels of SARS-CoV-2, podocyte markers (WT1, PODXL, NPHS1, NPHS2, MAFB), proximal tubular markers (SLC3A1 and SLC16A1), TMPRSS2 and NRP1. Data from a pool of 12 organoids/group is shown.

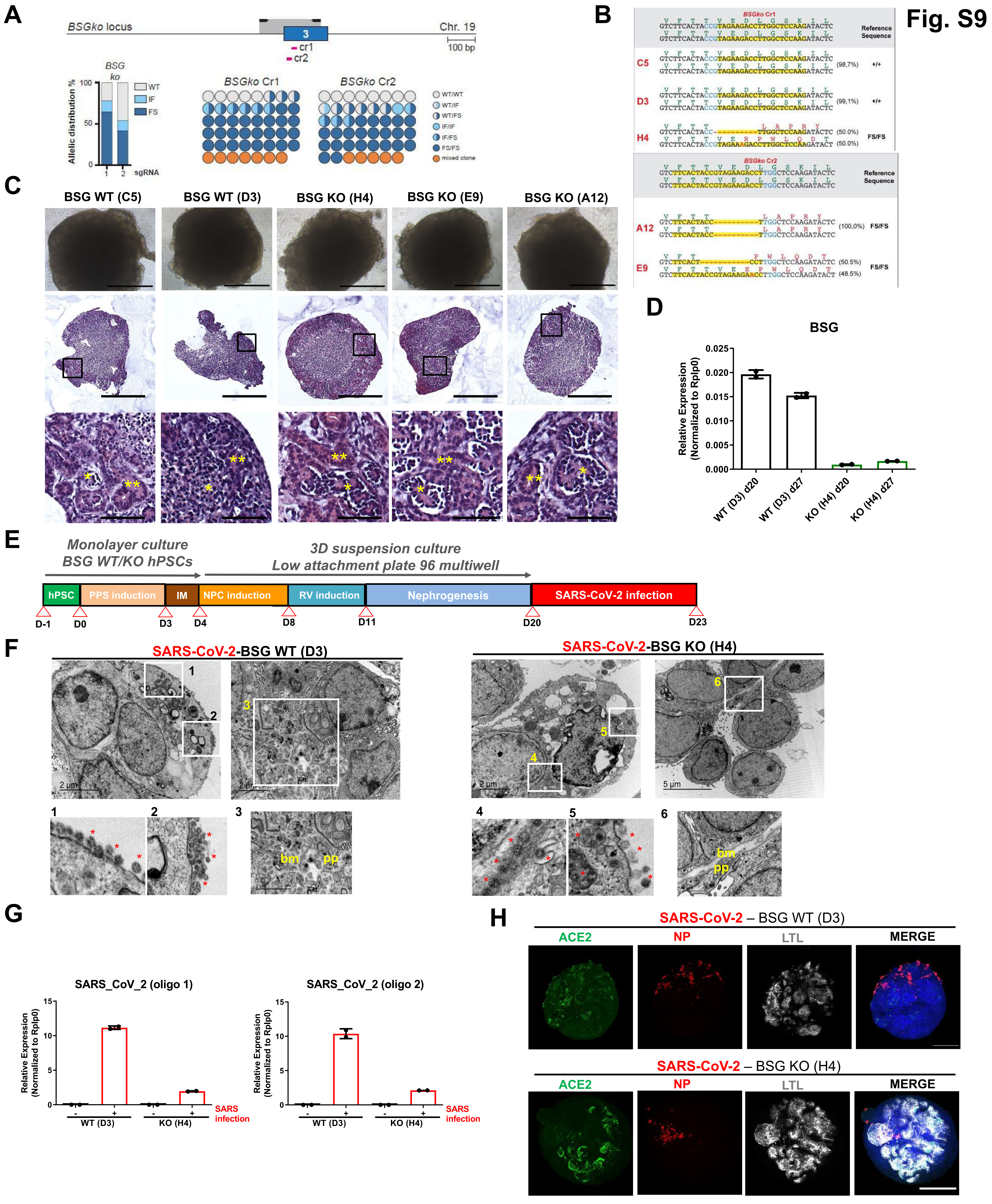

Fig. S9
A
BSGko locus
Chr. 19
3
cr1
cr2
100 bp
BSG ko
Allelic distribution %
100
50
0
1
2
sgRNA
WT
IF
FS
BSGko Cr1
BSGko Cr2
WT/WT
WT/IF
WT/FS
IF/IF
IF/FS
FS/FS
mixed clone
B
BSGko Cr1
Reference Sequence
C5
(98,7%)
+/+
D3
(99,1%)
+/+
H4
(50.0%)
(50.0%)
FS/FS
BSGko Cr2
Reference Sequence
A12
(100,0%)
FS/FS
E9
(50.5%)
(48.5%)
FS/FS
C
BSG WT (C5)
BSG WT (D3)
BSG KO (H4)
BSG KO (E9)
BSG KO (A12)
D
BSG
Relative Expression (Normalized to Rplp0)
0.025
0.020
0.015
0.010
0.005
0.000
WT (D3) d20
WT (D3) d27
KO (H4) d20
KO (H4) d27
E
Monolayer culture
BSG WT/KO hPSCs
3D suspension culture
Low attachment plate 96 multiwell
hPSC
PPS induction
IM
NPC induction
RV induction
Nephrogenesis
SARS-CoV-2 infection
D-1
D0
D3
D4
D8
D11
D20
D23
F
SARS-CoV-2-BSG WT (D3)
SARS-CoV-2-BSG KO (H4)
2 μm
5 μm
bm
pp
G
SARS_CoV_2 (oligo 1)
SARS_CoV_2 (oligo 2)
Relative Expression (Normalized to Rplp0)
15
10
5
0
WT (D3)
KO (H4)
SARS infection
H
SARS-CoV-2 – BSG WT (D3)
SARS-CoV-2 – BSG KO (H4)
ACE2
NP
LTL
MERGE

**Figure S9. SARS-CoV-2 Infection in BSG KO Human Kidney Organoids, Related to Figure 5.**

A. Schematic of Cas9/gRNA-targeting sites (pink arrows) in BSG *locus* showing exon structure (blue boxes) and PCR amplicons (light gray boxes in all figures in this study). Histogram shows allelic sequence distribution after the transfection of the different gRNAs in undifferentiated ES[4] cells expressing an inducible Cas9 (iCas9). wt, wild-type; mut, mutation; FS, frameshift.
B. Representative sequence of the wild type (+/+) or BSG mutant clones generated with the different gRNAs.
C. Representative bright field images of BSG WT and BSG KO kidney organoids. Scale bars, 250 µm. Same specimens were analyzed by Hematoxylin and Eosin staining showing tubular-like (**) and glomerular-like (*) structures. Scale bars, 250 µm, 50 µm (magnified views).
D. mRNA expression levels of BSG in BSG WT and BSG KO kidney organoids. Data from a pool of 12 organoids/group is shown.
E. Experimental scheme for the generation and viral infection of BSG WT and BSG KO kidney organoids.
F. TEM analysis of BSG WT and BSG KO kidney organoids infected with SARS-CoV-2 ($10^6$ virus particles/organoid as determined in Vero cells) and recovered at 3 dpi. Representative images of infected BSG WT specimens (left panel) show numerous viral particles in the cell surface of a dying cell (1,2). Details for podocyte-like cells (3) exhibiting podocyte related-structures including primary (pp) and the deposition of a basement membrane (bm). Scale bars, 2 µm; 200 nm (magnified views in 1 and 2); 2 µm (magnified views in 3). Representative images of infected BSG KO specimens (right panel) show numerous viral particles in the intercellular space (4) and in the cell surface (5). Details for podocyte-like cells (6) exhibiting podocyte related-structures including primary (pp) and the deposition of a basement membrane (bm). Scale bars, 2 µm and 5 µm, 200 nm (magnified views in 4 and 5); 1 µm (magnified view in 6).
G. Quantitative real time was performed at 3 dpi in Mock or SARS-CoV-2 infected ($10^6$ virus particles/organoid as determined in Vero cells) BSG WT and BSG KO specimens for the detection of SARS-CoV-2 mRNA expression levels. Data from a pool of 12 organoids/group is shown.
H. Immunofluorescence was performed at 3 dpi in Mock or SARS-CoV-2 infected ($10^6$ virus particles/organoid as determined in Vero cells) BSG WT and BSG KO kidney organoids. Representative confocal images show the detection of ACE2 (green), virus nuclear protein (NP, red), LTL (grey) and DAPI (blue). Scale bars, 250 µm.

**Figure S10**

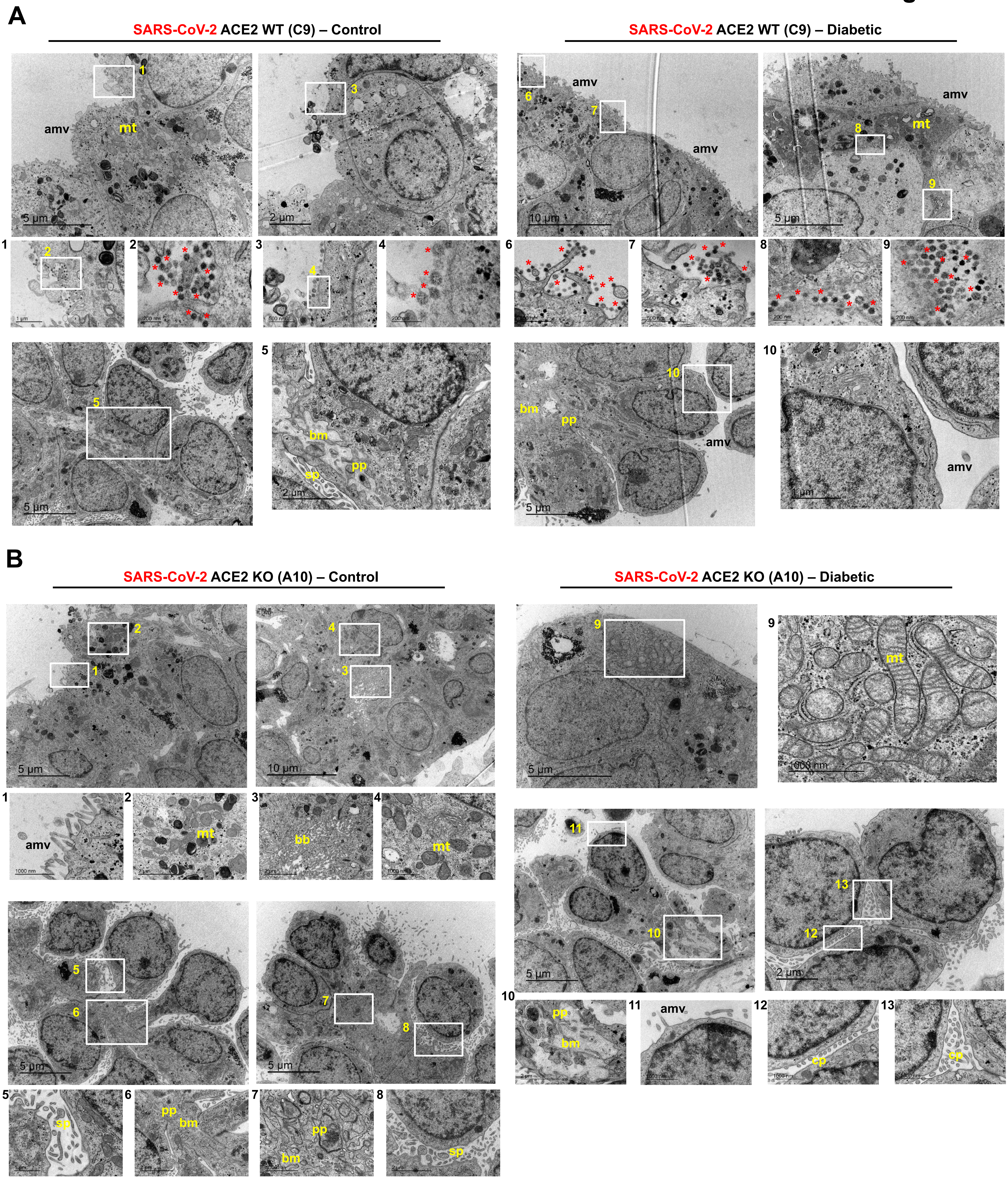
A
SARS-CoV-2 ACE2 WT (C9) – Control
SARS-CoV-2 ACE2 WT (C9) – Diabetic
amv
mt
bm
pp
sp
5 µm
2 µm
10 µm
1 µm
B
SARS-CoV-2 ACE2 KO (A10) – Control
SARS-CoV-2 ACE2 KO (A10) – Diabetic
amv
mt
bb
pp
bm
sp
cp
5 µm
10 µm
2 µm
1000 nm

**Figure S10. TEM Analysis of SARS-CoV-2 Infected ACE2 WT and ACE2 KO Human Kidney Organoids, Related to Figure 6.**

A. TEM analysis of ACE2 WT (C9) kidney organoids exposed to Control or Diabetic conditions infected with SARS-CoV-2 ($10^6$ virus particles/organoid as determined in Vero cells) and recovered at 1 dpi. Representative images of infected ACE2 WT (C9) Control specimen show multiple viral particles (asterisks) in contact with the apical microvilli (amv) of tubular-like cells (1-4). Details for podocyte-like cells exhibiting podocyte related-structures including primary processes (pp) (5) and the deposition of a basement membrane (bm) (5) are shown. Scale bars, 5 μm, 2 μm; 1 μm (1); 200 nm (2); 500 nm (3); 200 nm (4); 2 μm (5). Representative images of infected ACE2 WT (C9) Diabetic specimen show numerous viral particles (asterisks) in contact with the apical microvilli (amv) of tubular-like cells (6,7) and in-between tubular-like cells in close contact with cell membranes (8,9). Podocyte-like cells with apical microvilli (amv), primary cell processes (pp) and basal deposition of a basement membrane are also shown (10). Scale bars, 10 μm, 5 μm; 500 nm (6,7); 200 nm (8,9); 1 μm (10).

B. TEM analysis of ACE2 KO (A10) kidney organoids exposed to Control or Diabetic conditions infected with SARS-CoV-2 ($10^6$ virus particles/organoid as determined in Vero cells) and recovered at 1 dpi. Representative images of infected ACE2 KO (A10) Control specimen show tubular-like cells with apical microvilli (amv), high mitochondrial content (mt) and dense brush borders (bb) (1-4). Glomerular-like structures with podocyte-like cells that exhibit primary (pp) and secondary (sp) cell processes, and deposition of basement membrane (bm) are shown (5-8). Scale bars, 5 μm, 10 μm; 1 μm (1,4,5); 2 μm (2,3,6,8). Representative images of infected ACE2 KO (A10) Diabetic specimen show tubular-like cells with high mitochondrial content (mt) (9). Podocyte-like cells display primary cell processes (pp), deposition of basement membrane (bm) and apical microvilli (amv) (10-13). Scale bars, 5 μm, 2 μm; 1 μm (9,11-13); 2 μm (10).

#### Extended data 1 -Related to Figure 1

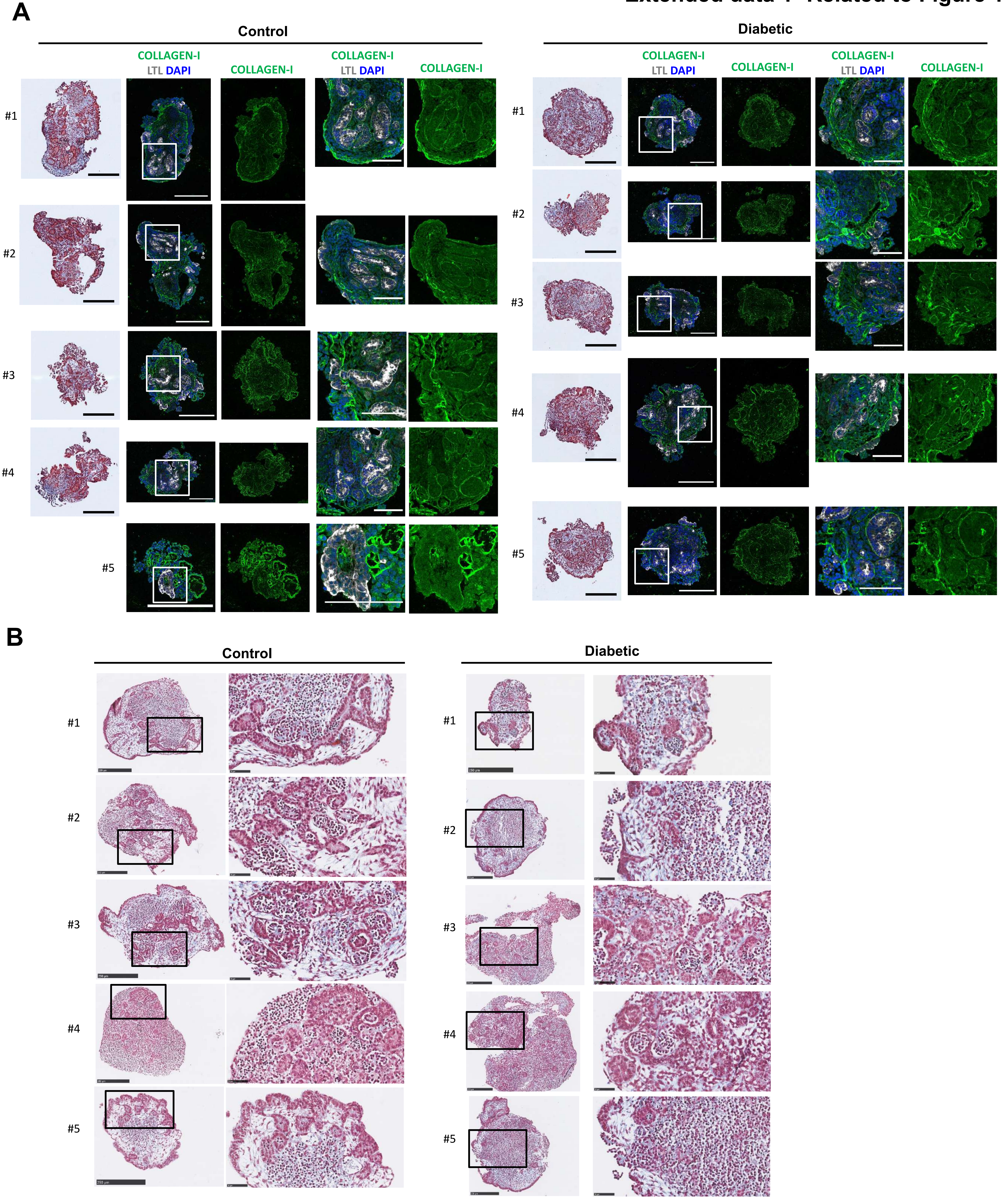

**Extended Data 1, Related to Figure 1.**

A. Representative Trichrome Masson staining in kidney organoids exposed to Control or Diabetic conditions for 7 days. Scale bars, 250 µm. Consecutive sections were stained for COLLAGEN-I (green), LTL (grey) and DAPI (blue). Scale bars, 250 µm, 100 µm (magnified views). *n*= *5* organoids/group.

B. Trichrome Masson staining in Control and Diabetic kidney organoids. Scale bars, 250 µm, 50 µm (magnified views). *n*= *5* organoids/group.

### Extended data 2 -Related to Figure 1

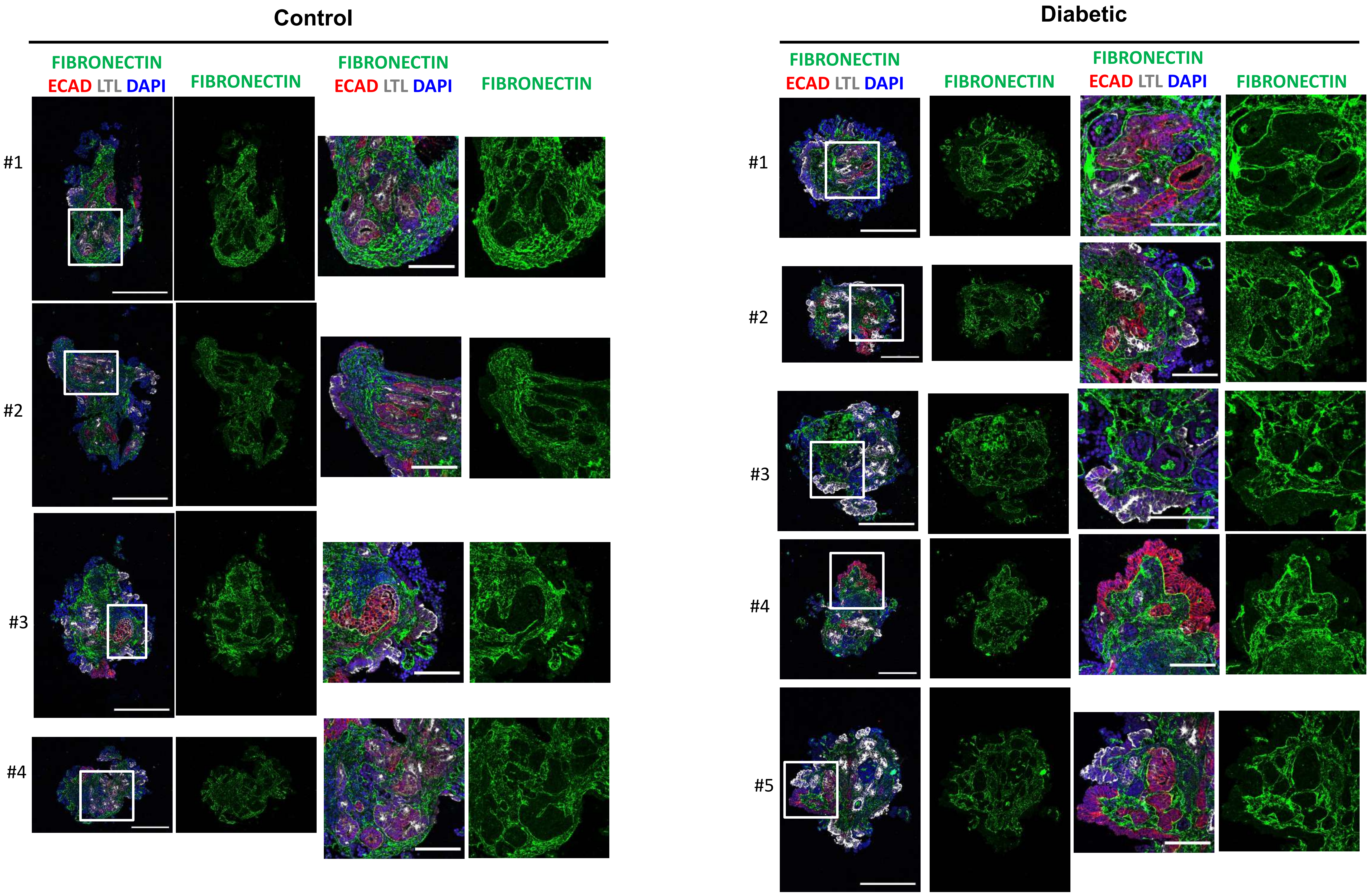

**Extended Data 2, Related to Figure 1.**

Representative immunofluorescence staining of FIBRONECTIN (green), ECADHERIN (ECAD; red), LTL (grey) and DAPI (blue) exposed to Control or Diabetic conditions for 7 days. Scale bars, 250 μm, 100 μm (magnified views). *n= 4* Control organoids*; n=5* Diabetic organoids.

#### Extended Data 3 -Related to Figure 1

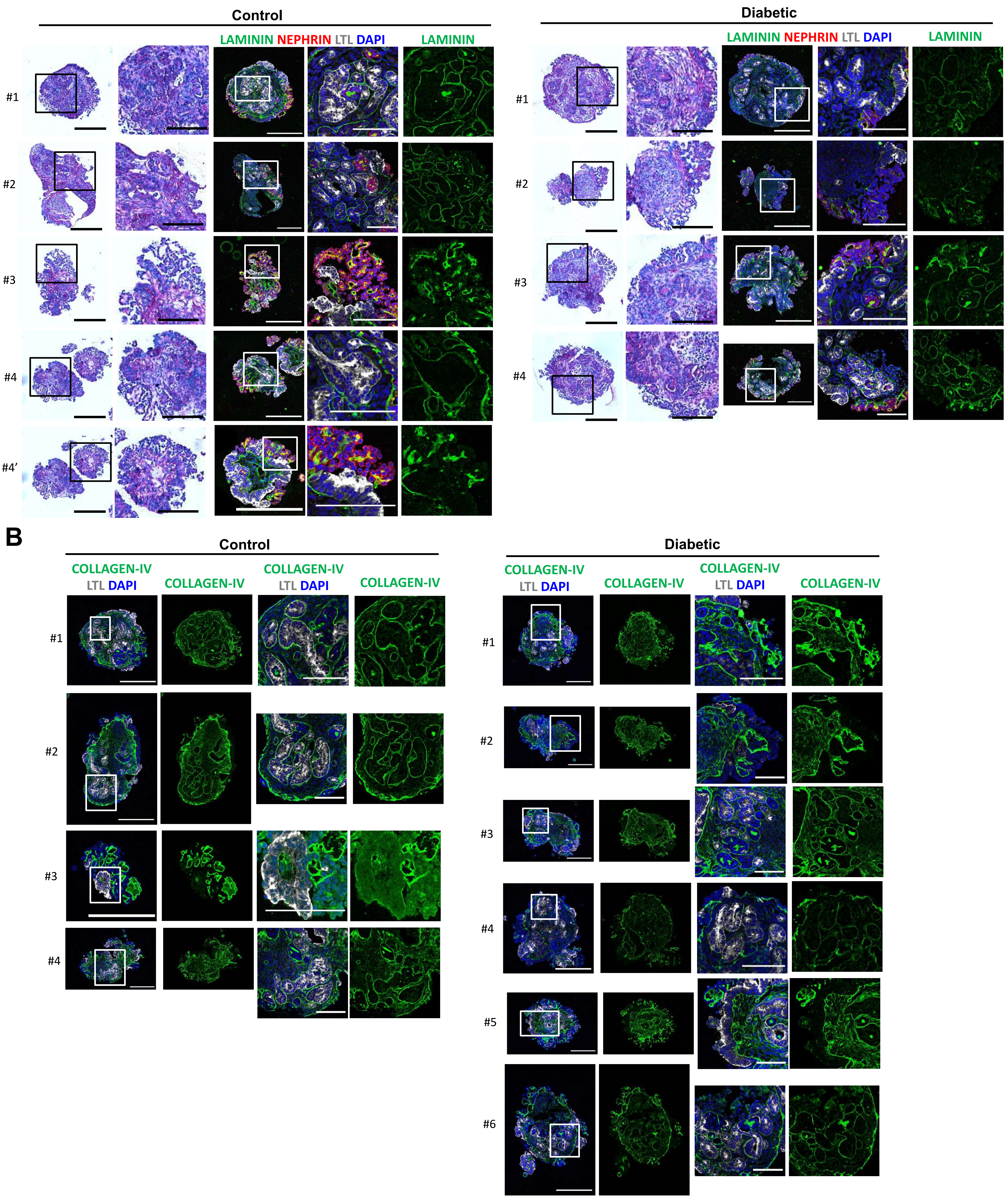

**Extended Data 3, Related to Figure 1.**

A. Representative Periodic Acid-Schiff (PAS) staining in kidney organoids exposed to Control or Diabetic conditions for 7 days. Scale bars, 250 μm, 100 μm (magnified views). Consecutive sections were stained for LAMININ (green), NEPHRIN (red), LTL (grey) and DAPI (blue). Scale bars, 250 μm, 100 μm (magnified views). *n= 4* organoids/group.

B. Representative immunofluorescence staining of COLLAGEN-IV (green), LTL (grey) and DAPI (blue) in kidney organoids exposed to Control or Diabetic conditions for 7 days. Scale bars, 250 μm, 100 μm (magnified views). *n= 4* Control organoids*; n=6* Diabetic organoids.

### Extended data 4 -Related to Figure 1

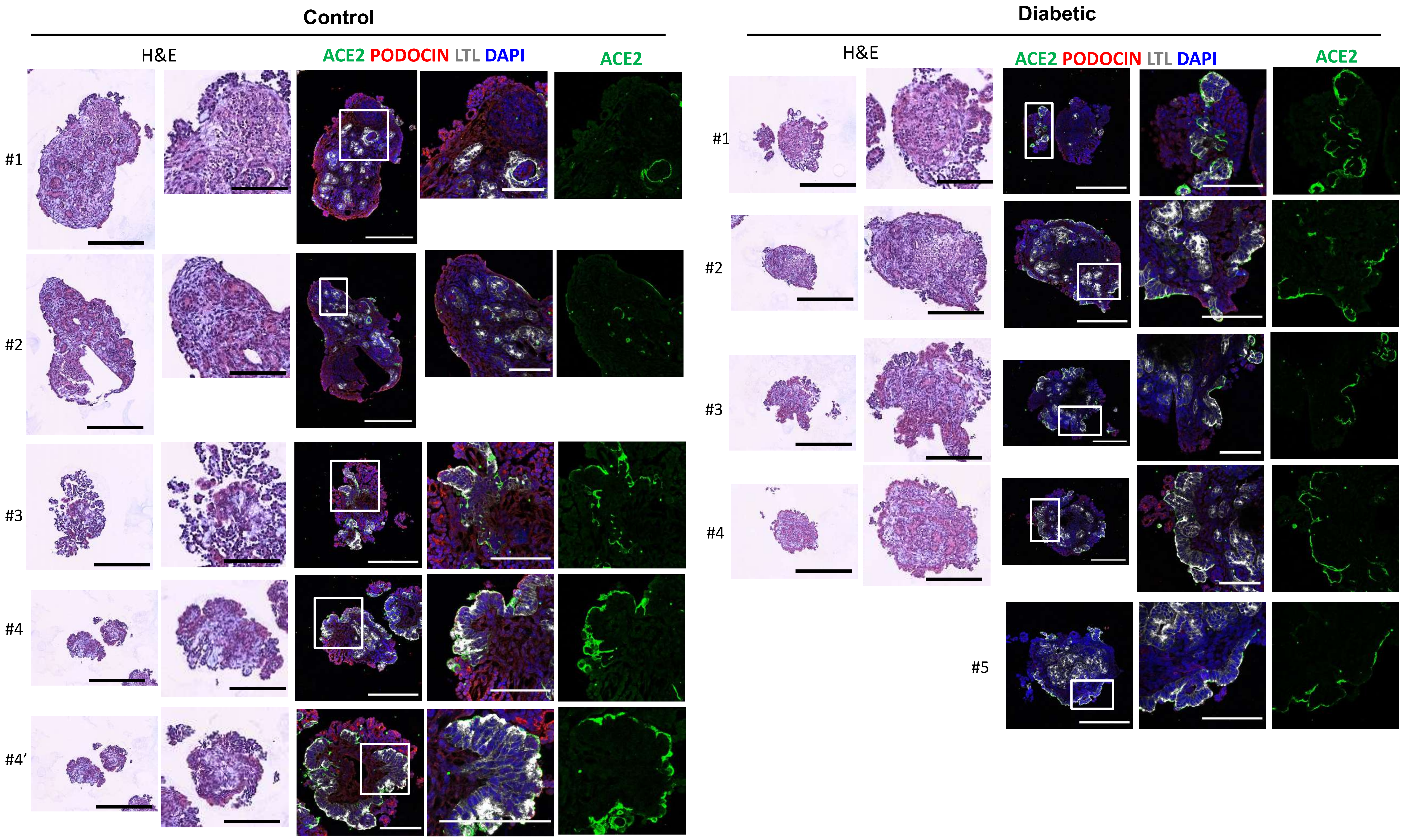

**Extended Data 4, Related to Figure 2.**

Representative Hematoxylin and Eosin (HE) staining in kidney organoids exposed to Control or Diabetic conditions for 7 days. Scale bars, 250 µm, 100 µm (magnified views). Consecutive sections were stained for ACE2 (green), PODOCIN (red), LTL (grey) and DAPI (blue). Scale bars, 250 µm, 100 µm (magnified views). *n= 4* Control organoids*; n=5* Diabetic organoids.
